# Pax Protein Depletion in Proximal Tubules Triggers Conserved Mechanisms of Resistance to Acute Ischemic Kidney Injury and Prevents Transition to Chronic Kidney Disease

**DOI:** 10.1101/2023.10.03.559511

**Authors:** Jeffrey A. Beamish, Asha C. Telang, Madison C. McElliott, Anas Al-Suraimi, Mahboob Chowdhury, Jenna T. Ference-Salo, Edgar A. Otto, Rajasree Menon, Abdul Soofi, Joel M. Weinberg, Sanjeevkumar R. Patel, Gregory R. Dressler

**Author notes:** **CORRESPONDENCE:** Jeffrey A. Beamish, Division of Nephrology, Department of Internal Medicine School of Medicine, University of Michigan, Taubman Center, 3^rd^ Floor, Reception C 1500 E. Medical Center Dr., SPC 5364 Ann Arbor, MI 48109-5364. **SOURCES OF SUPPORT** University of Michigan O’Brien Kidney Center (DK-P30-081943 to J.A.B.), NIH K08 DK125776 (to J.A.B.), NIH R01 DK054740 to (G.R.D.), NIH R01 DK073722 to (G.R.D.), A gift from Josh and Audrey Rumsey in memory of Isaiah.

## Abstract

Acute kidney injury (AKI) is a common condition that lacks effective treatments. In part this shortcoming is due to an incomplete understanding of the genetic mechanisms that control pathogenesis and recovery. Pax2 and Pax8 are homologous transcription factors with overlapping functions that are critical for kidney development and are re-activated in AKI. In this report, we examined the role of Pax2 and Pax8 in recovery from ischemic AKI. We found that Pax2 and Pax8 are upregulated after severe AKI and correlate with chronic injury. Surprisingly, we then discovered that proximal-tubule-selective deletion of Pax2 and Pax8 resulted in a less severe chronic injury phenotype. This effect was mediated by protection against the acute insult, similar to preconditioning. Prior to injury, Pax2 and Pax8 mutant mice develop a unique subpopulation of S3 proximal tubule cells that display features usually seen only in acute or chronic injury. The expression signature of these cells was strongly enriched with genes associated with other mechanisms of protection against ischemic AKI including caloric restriction, hypoxic preconditioning, and female sex. Taken together, our results identify a novel role for Pax2 and Pax8 in mature proximal tubules that regulates critical genes and pathways involved in both injury response and protection from ischemic AKI.

**TRANSLATIONAL STATEMENT:** Identifying the molecular and genetic regulators unique to the nephron that dictate vulnerability to injury and regenerative potential could lead to new therapeutic targets to treat ischemic kidney injury. Pax2 and Pax8 are two homologous nephron-specific transcription factors that are critical for kidney development and physiology. Here we report that proximal-tubule-selective depletion of Pax2 and Pax8 protects against both acute and chronic injury and induces an expression profile in the S3 proximal tubule with common features shared among diverse conditions that protect against ischemia. These findings highlight a new role for Pax proteins as potential therapeutic targets to treat AKI.

## INTRODUCTION

Ischemic acute kidney injury (AKI) is a common condition associated with significant morbidity, mortality, and cost^1,2^. Remarkably, proximal tubular epithelial cells can regenerate after injury, restoring some or all kidney function^3,4^. However, regeneration is rarely complete. Some tubular epithelial cells remain in a non-physiological state and may drive a transition from AKI to chronic kidney disease (CKD) and fibrosis for many patients^5–8^. Thus, defining the molecular and genetic mechanisms that determine the response to injury and the success of regeneration is critical for developing effective treatments for AKI and its sequelae, which have been elusive in the clinical setting.

Successful regeneration after AKI requires de-differentiation and proliferation of surviving epithelial cells followed by re-differentiation back to mature kidney epithelium. Elements of this process resemble kidney development^9,10^, suggesting that regenerating cells may co-opt developmental programs. Pax2 and Pax8 are two homologous proteins that are critical for kidney development and that are upregulated in proximal tubule cells during AKI and recovery^11,12^. Pax2 may facilitate proliferation after injury and both Pax2 and Pax8 remain upregulated in chronically injured proximal tubules^6,13–15^.

Pax2 and Pax8, along with Pax5 which is not expressed in the kidney, are a family of highly homologous proteins with identical DNA binding domains and significant homology in their carboxy-terminal octapeptide, homeodomain, and activation domain sequences^16^. At the molecular level, Pax2 and Pax8 proteins interact with PTIP to recruit a KMT2C/D histone methyl-transferase complex and maintain chromatin accessibility by depositing histone H3 lysine 4 methylation marks^17^. However, Pax2/8 can also silence gene expression by interacting with the Grg/TLe family of co-repressors to recruit a Polycomb repressor 2 complex^18^. In development, Pax2 is essential for the earliest phase of kidney induction within the intermediate mesoderm and continues to be expressed in the nephron progenitor cells as they proliferate. However, mature nephrons down-regulate Pax2 but continue to express Pax8 into adulthood.^19,20^. In the adult, Pax2 and Pax8 are most strongly expressed in the collecting tubules and medulla where they regulate transporters essential for normal water and solute homeostasis in a redundant fashion^21^. Despite evidence they play a key role in nephron development and are reactivated upon injury, the function of Pax2 and Pax8 in AKI has not been clearly defined.

In this report, we examined the role of Pax2 and Pax8 in the response to ischemic kidney injury. Because of their redundant functions in development and physiology, we generated mice with conditional deletions of both Pax2 and Pax8 in proximal tubules of the post-natal kidney. These mice were subjected to ischemia-reperfusion injury (IRI) to model ischemic acute tubular injury, the most common form of severe AKI seen clinically^22,23^. Surprisingly, these Pax2 and Pax8 double mutant mice showed enhanced regeneration after IRI and decreased progression to chronic injury. These effects were mediated by pre-injury gene expression changes localized to the S3 proximal tubule that matched the protective transcription signature of female sex, caloric restriction, and hypoxic preconditioning, producing striking global protection of renal function. These findings highlight critical conserved pathways of protection against ischemic AKI and suggest a novel role for Pax2 and Pax8 in ischemic stress.

## METHODS

### Animals

All studies involving mice were approved by the Institutional Animal Care & Use Committee (IACUC) at the University of Michigan and performed in accordance with the NIH Guide for the Care and Use of Laboratory Animals. Strains were derived from phosphoenolpyruvate carboxykinase (PEPCK) Cre mice^24,25^, GFP Cre reporter mice (Gt(Rosa)26Sor^tm4^(ACTB–tdTomato,–EGFP)^luo^, Jackson Labs, Stock No. 007676), and mice with floxed Pax2 and Pax8 alleles^21^. Breeding details and strain genetics are outlined in the Supplementary Methods and Supplementary Figure 1.

### Ischemia-reperfusion injury (IRI)

Two surgical models were used. Severe unilateral IRI was induced by clamping the left renal pedicle for 30 min with a constant animal body temperature of 38 ⁰C without contralateral nephrectomy. For moderate unilateral ischemia, a right nephrectomy was performed prior to clamping the left renal pedicle for 25 min with a constant animal body temperature of 37 ⁰C. Detailed surgical procedures are provided in the Supplementary Methods.

### Sample preparation

After euthanasia with carbon dioxide, blood was collected for serum and explanted kidney samples were divided in symmetric pieces then either flash frozen or fixed in 4% paraformaldehyde and paraffin embedded. Nuclei, protein, and RNA were extracted from frozen kidney pieces. Detailed procedures are outlined in the Supplementary Methods.

### Single nucleus preparation and RNA seq

Nuclei were isolated from flash frozen whole kidney samples from uninjured mice then fixed and cDNA libraries generated using an "Evercode™ WT Mega v2" plate (Parse Biosciences) per manufacturer protocol. Count matrices were generated from sequencing data using the Parse Biosciences data analysis pipeline and read into the Seurat R package^26^ (v4.3.0) to cluster, visualize, and analyze the dataset. Detailed analysis procedures are provided in the Supplementary Methods.

### Statistics

Data are presented as mean and error bars represent standard deviation unless otherwise specified. Statistical analysis was performed using the test indicated.

Additional details for all methods are provided in the Supplementary Methods.

## RESULTS

### Pax2 and Pax8 protein increases in proximal tubules after severe AKI and is associated with incomplete repair

We examined Pax2 and Pax8 expression in proximal tubules after injury using transgenic mice with the phosphoenolpyruvate carboxy kinase (PEPCK) Cre driver to activate a GFP reporter allele. These animals, with wildtype Pax2 and Pax8 loci, PEPCK-Cre, and a GFP reporter, are denoted WT in this report. This approach, after embedding samples in paraffin to extinguish native GFP and tdTomato fluorescence, allowed us to costain GFP-marked Cre activity and targets of interest. WT animals expressed GFP in the proximal tubules with high specificity and with substantial but heterogeneous labeling of the S1 and S2 segments and near-complete labeling of the S3 segment at the corticomedullary junction (Figure 1a). We next subjected these animals to severe unilateral IRI, with preservation of the contralateral kidney, to model the AKI-to-CKD transition (Figure 1b). We used deep-learning-assisted image segmentation^27^ to classify nuclei of GFP+ proximal tubule cells according to co-staining for Vcam1, KIM-1, Pax2, or Pax8 across the entire cortex and outer stripe of the outer medulla (OSOM) 14 d after injury using immunostaining (example analysis is shown in Supplementary Figure S2). This time corresponds to a peak population of cells with a persistent injury phenotype marked by Vcam1 and KIM-1^6^. We observed the expected increase in Vcam1+ and KIM-1+ in GFP+ proximal tubule cells in the injured kidneys relative to the uninjured contralateral kidneys (Figure 1c). We also noted a significant increase in the number of Pax2+ and Pax8+ nuclei (Figure 1d) that was strongly correlated, between animals, with Vcam1 (Figure 1e). Pax8 was correlated with KIM-1 as well (Supplementary Figure S3). These results indicate that Pax2 and Pax8 are upregulated in proximal tubules in chronic injury states after AKI.

**Figure 1.**
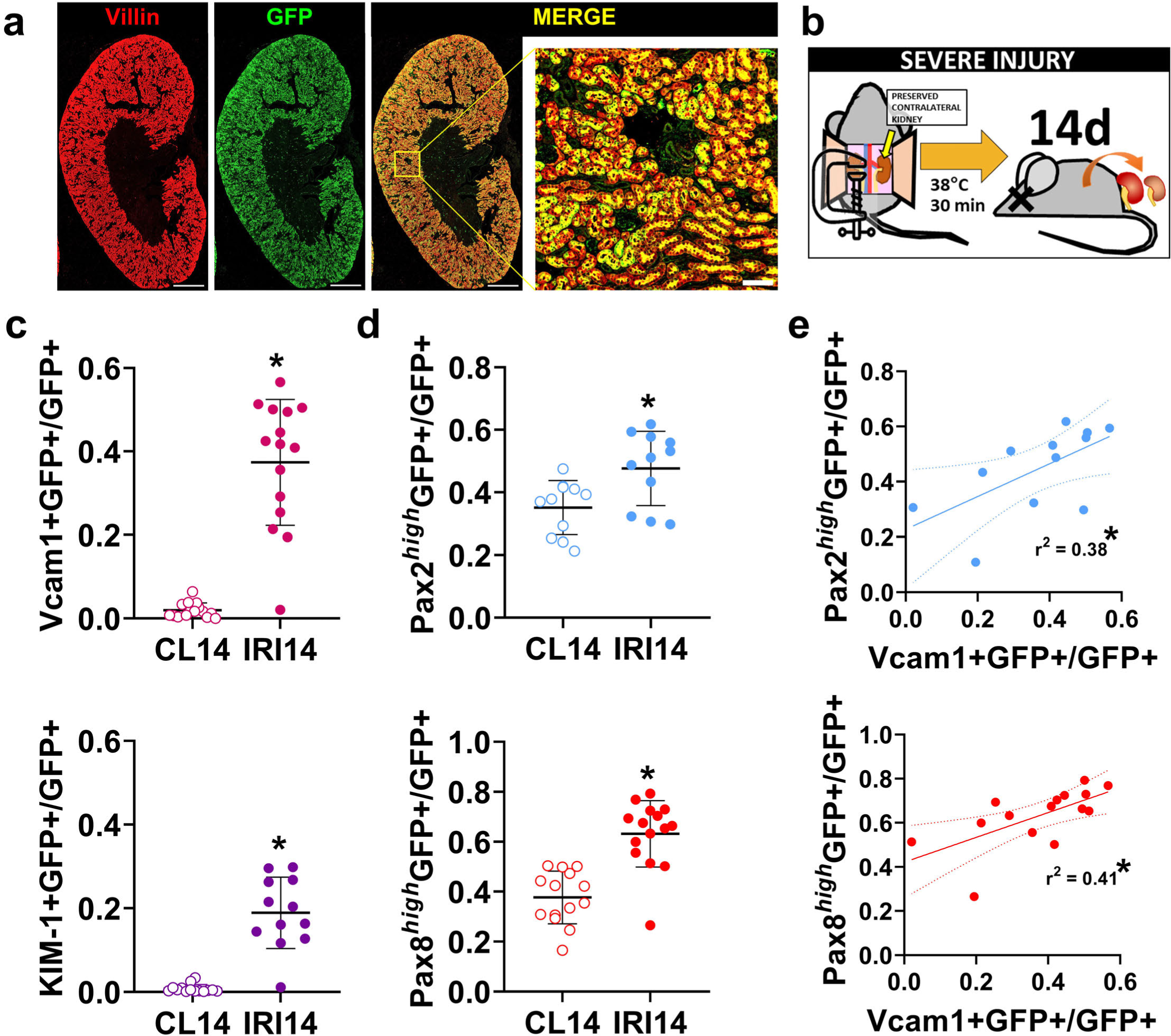
Pax2 and Pax8 protein increases in proximal tubules after severe AKI and correlates with incomplete recovery. (**a**) Immunofluorescence staining of an uninjured kidney from a transgenic mouse with stable GFP expression (green) induced by phosphoenolpyruvate carboxykinase (PEPCK) Cre and used to identify proximal tubules. Villin (red) also marks mature proximal tubules. Scale bars: 1 mm, 100 μm (inset). (**b**) Animals were subjected to severe ischemia reperfusion injury (IRI) using a 30 min clamp time at a constant body temperature of 38 °C. The contralateral kidney was preserved. (**c**) The fraction of GFP+ proximal tubule cells co-expressing Vcam1 and KIM-1, measured using deep-learning assisted tissue cytometry (Supplementary Figure S2). Each point represents analysis of all nuclei in the cortex + outer stripe of the outer medulla (OSOM) from a single section from each animal (*: *P* < 0.0001, paired t-test, *N =* 11-15 animals). (**d**) The fraction of GFP+ proximal tubule cells with strong staining for Pax2 (Pax2^high^) or Pax8 (Pax8^high^) measured as in (**c**). (**e**) Correlation of Pax2 or Pax8 with Vcam1 expression in GFP+ proximal tubule cells. Each point represents measurements from the same animal but from separate sections.

### Mutant Pax2 and Pax8 proximal tubule cells are more likely to undergo successful repair after IRI

Because both Pax2 and Pax8 expression were correlated with chronic injury, we reasoned that Pax2 and Pax8 may contribute to incomplete repair. To test this hypothesis, we replaced both wildtype Pax2 and Pax8 with conditional floxed alleles and deleted both with the PEPCK-Cre driver in the proximal tubule (detailed genetics in Supplementary Figure S1, hereafter referred to as mutants). Mutant mice showed significant depletion of both Pax2 and Pax8 in proximal tubules, marked by GFP (Figure 2, Supplementary Figure S4). Pax2 and Pax8 depletion was strongly localized to S3, where most GFP+ cells contained no detectable Pax2 or Pax8 protein. (Supplementary Figure S5). Both WT and mutant strains had a similar baseline histology, BUN, and creatinine (Supplementary Figure S6). We then performed severe unilateral ischemia reperfusion injury on WT control and mutant mice and assessed recovery after 14 d. Mutant mice showed less chronic injury histologically (Figure 3a). Collagen I α1, Vcam1, and KIM-1 protein were less abundant in mutant compared with WT kidneys (Figure 3b). Likewise, we found less Vcam1+ and KIM-1+ tubules in the cortex + OSOM of mutants, whereas mature proximal tubules, marked by villin, were more abundant in the mutants (Figure 3c-d). We next asked whether Pax2 and Pax8 depleted mutant cells were less likely to show evidence of incomplete repair. Within this population, marked by Vcam1 or KIM-1, we measured significantly fewer GFP+ cells in the mutant animals (Figure 4, Supplementary Figure S7). To confirm these changes were associated with depletion of Pax2 and Pax8, we measured Pax2 and Pax8 staining in GFP+ cells, subdivided by KIM-1 co-staining. We found that GFP+, KIM-1-negative cells were disproportionately depleted of Pax2 and Pax8 (Supplementary Figure S7), supporting the conclusion that loss of Pax2 and Pax8 prevents development of a chronic injury phenotype.

**Figure 2.**
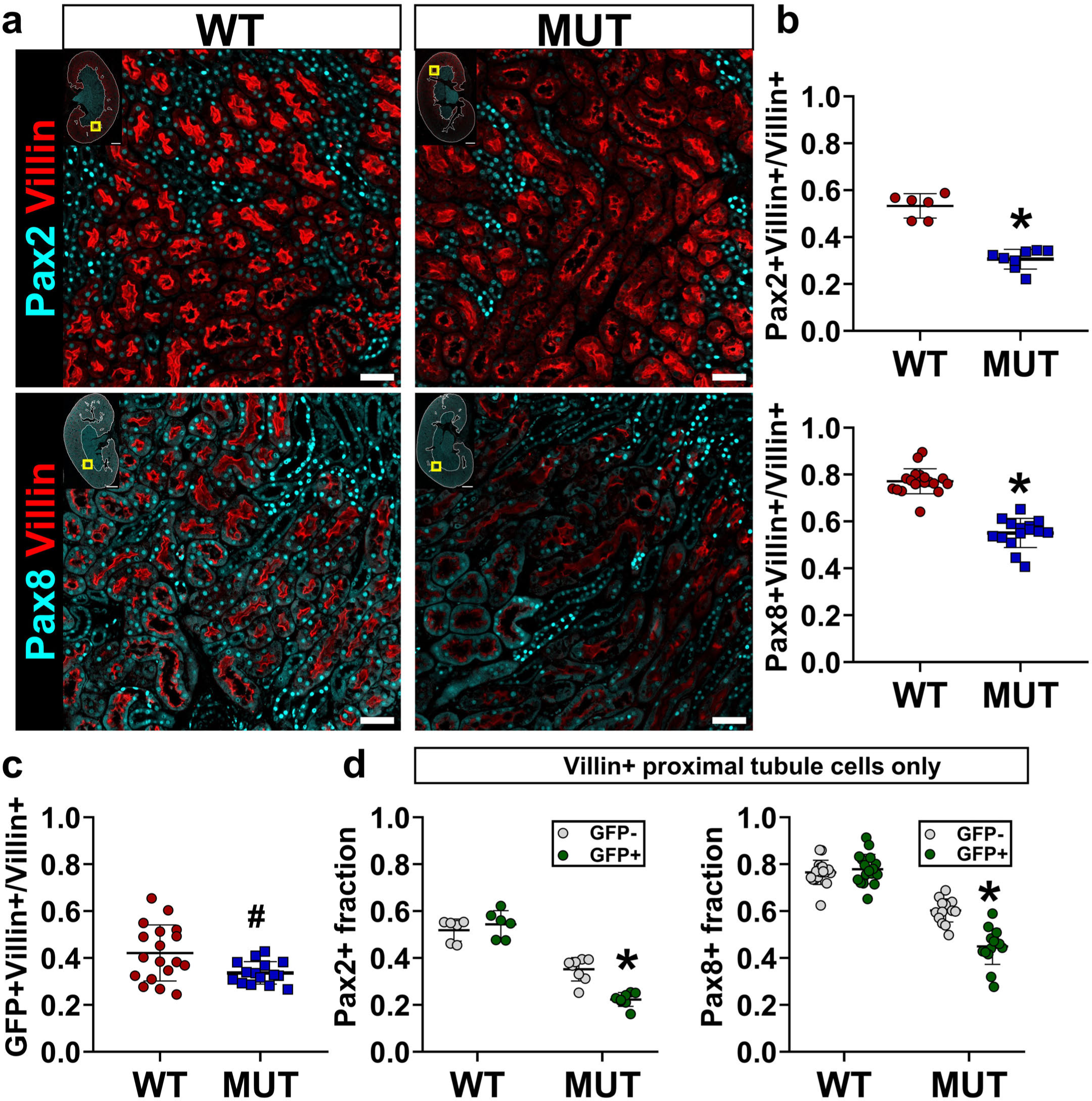
Pax2 and Pax8 proximal tubule mutants show significant depletion of Pax2 and Pax8 protein, marked by GFP. (**a**) Representative images showing Pax2 (top, cyan) and Pax8 (bottom, cyan) co-stained for the proximal tubule marker villin (red). Yellow box in inset shows image location in the larger section, white outline shows the cortex + OSOM used for analysis. Corresponding images showing Pax2 and Pax8 with GFP are provided in Supplementary Figure S4 and Supplementary Figure S5. Scale bars: 1 mm (full section), 50 μm (region of interest). (**b**) Deep-learning-assisted quantification of the fraction of cells expressing Pax2 and Pax8 among villin+ proximal tubule cells (*: *P* < 0.0001, t-test, *N =* 6-18 animals). (**c**) Deep-learning-assisted quantification of the fraction of GFP+ cells in the population of villin+ proximal tubule cells (#: *P* = 0.01, t-test, *N =* 15-18 animals). (**d**) Quantification of the Pax2+ and Pax8+ expression in subpopulations of villin+ proximal tubule cells with and without GFP expression (*: *P* < 0.0001 compared with all other groups, two-way ANOVA with Sidak’s post-hoc test, *N =* 6-18 animals). Strain abbreviations-WT: Pax2^wt/wt^, Pax8^wt/wt^, PEPCK Cre^+/y^, mT/mG^+/+^; MUT: Pax2^fl/fl^, Pax8^fl/fl^, PEPCK Cre^+/y^, mT/mG^+/+^.

**Figure 3.**
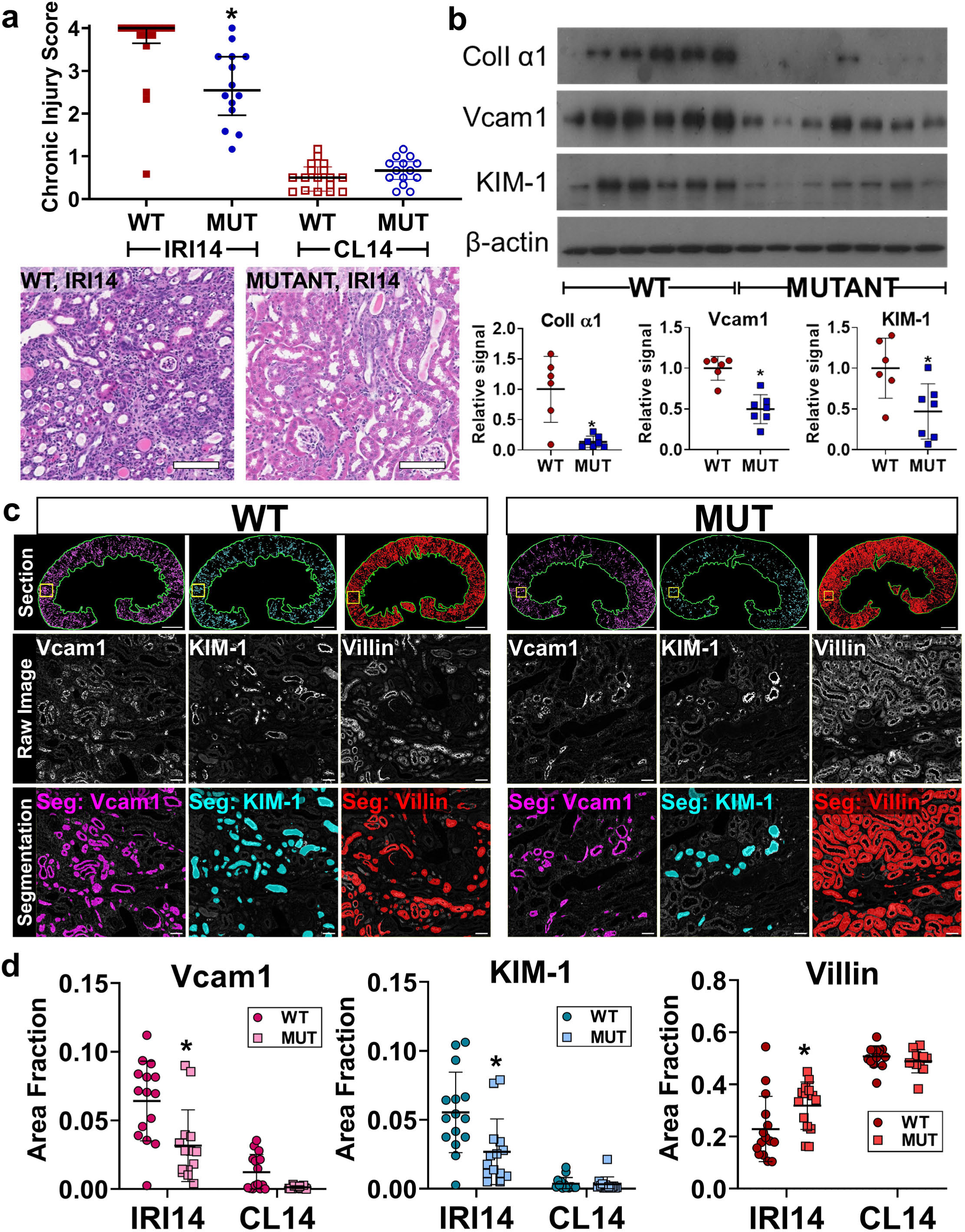
Proximal tubule Pax2 and Pax8 depletion protects against chronic injury. (**a**) Chronic injury score of hematoxylin and eosin stained sections from injured (IRI14) and contralateral uninjured (CL14) kidneys 14 d after IRI (median + interquartile range, *: *P* = 0.002, Mann-Whitney test, *N =* 14-16 animals). Scale bar: 100 µm. (**b**) Western blot of whole kidney protein lysates from IRI14 samples and associated densitometry analyses normalized to β-actin (*: *P* = 0.01, 0.002, 0.02 for Collagen I α1, Vcam1, and KIM-1 respectively, t-test, *N =* 6-7 animals). (**c**) Representative examples of segmentation of Vcam1, KIM-1, and villin. Analysis was performed over the entire cortex + OSOM for each animal (green outline, top), the raw grayscale image is shown for each stain accompanied by representative segmentations (color overlay). Scale bars: 1 mm (full section), 50 µm (inset). (**d**) Quantification of segmentation area in the cortex + OSOM for Vcam1, KIM-1, and villin (*: p= 0.0002, 0.0004, 0.04, respectively, two-way ANOVA with Sidak’s post-hoc test, *N =* 14-15 animals). Strain abbreviations-WT: Pax2^wt/wt^, Pax8^wt/wt^, PEPCK Cre^+/y^, mT/mG^+/+^; MUT: Pax2^fl/fl^, Pax8^fl/fl^, PEPCK Cre^+/y^, mT/mG^+/+^.

**Figure 4.**
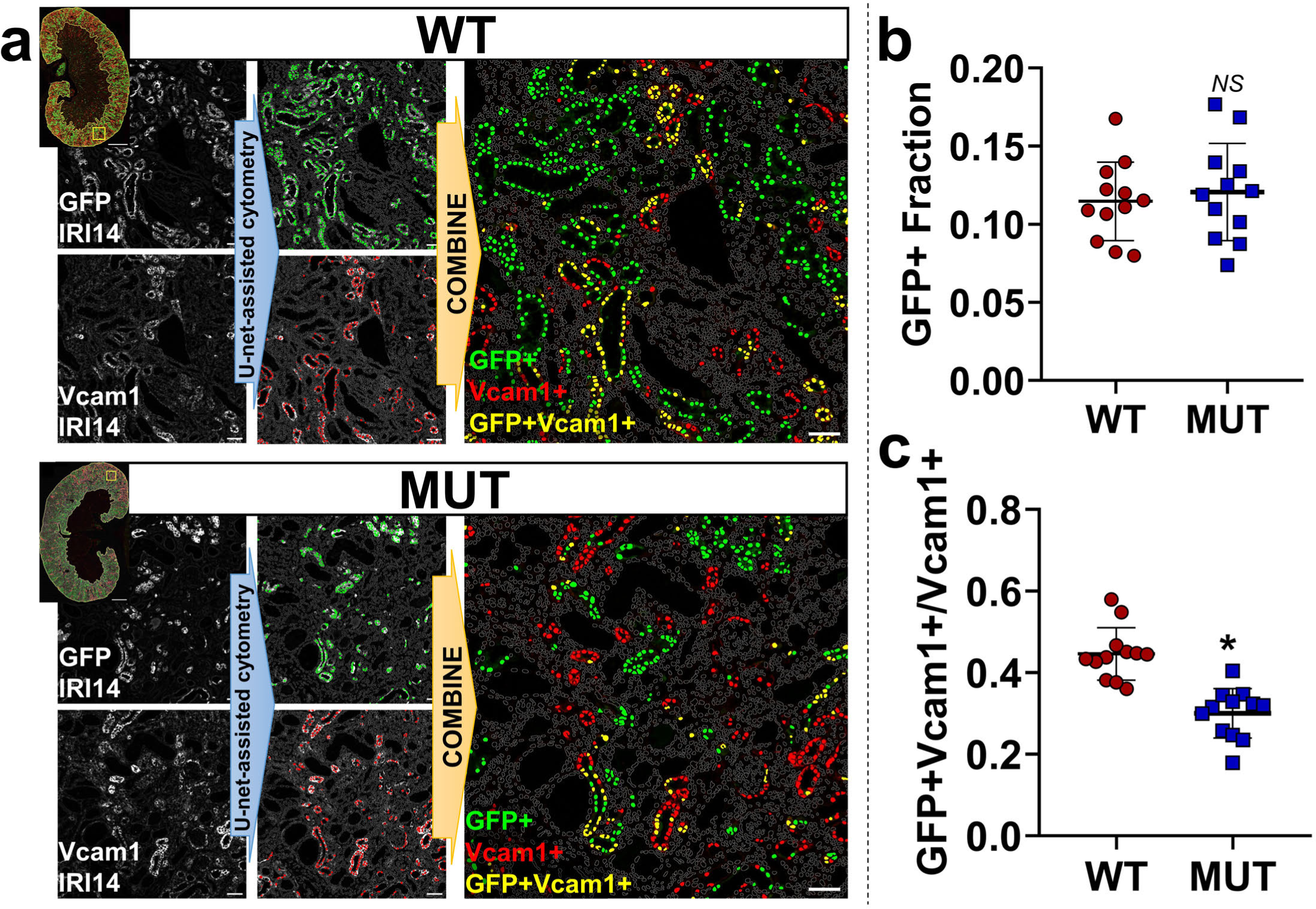
Pax2 and Pax8 mutant cells are less likely to contribute to incomplete repair. (**a**) Representative images showing U-Net deep-learning-assisted cytometry of sections from animals 14 d after IRI (IRI14). Grayscale images (left) were segmented and used to classify positive cells (defined by their nuclei, marked in color). Positive co-stained cells were determined by the intersection of the two sets (yellow, right). Gray outlines represent nuclei negative for both markers. Scale bars: 1 mm (full section), 50 μm (region of interest). (**b**) Quantification of the overall GFP+ fraction, which marks wild type or Pax2 and Pax8 depleted proximal tubule cells in WT and MUT strains respectively (NS: *P* = 0.61, t-test, *N =* 12 animals). (**c**) Quantification of the fraction of GFP+ cells in the subpopulation of Vcam1+ cells which marks incomplete repair. Measurements were made across the cortex + OSOM for each section (*: *P* < 0.0001, t-test, *N =* 12 animals). Strain abbreviations-WT: Pax2^wt/wt^, Pax8^wt/wt^, PEPCK Cre^+/y^, mT/mG^+/+^; MUT: Pax2^fl/fl^, Pax8^fl/fl^, PEPCK Cre^+/y^, mT/mG^+/+^.

To control for the possibility of segregation of unrelated protective traits, we also generated colonies of animals with alternative genetics where Pax mutation was controlled by the presence or absence of a PEPCK-Cre driver and all mice were homozygous for floxed Pax2 and Pax8 alleles (Supplementary Figure S8). After injury, these alternative mutant animals likewise showed fewer Vcam1+ and KIM-1+ tubules and expressed less Collagen I α1 protein (Supplementary Figure S8). Together these findings indicate that depletion of Pax2 and Pax8 in the proximal tubule results in less chronic injury after severe IRI because mutant cells are less likely to develop an incomplete repair phenotype.

### Proximal tubule Pax2 and Pax8 loss protects against ischemic injury and promotes early repair

Incomplete repair can result either from defective repair processes or from more severe injury. To evaluate the latter possibility, we measured responses in the acute phase, 2 d after IRI. We changed to a moderate injury model with contralateral nephrectomy to facilitate assessment of kidney function. We observed that mutant mice developed less severe AKI as measured by BUN (Figure 5a) and acute histological injury scores (Figure 5b). We also detected lower expression of acute injury markers KIM-1 and Krt20 in bulk RNA from mutant kidneys (Figure 5c). Mutant mice also showed a more robust and consistent proliferative response marked by Ki67 (Figure 5d). Furthermore, the cell population expressing the early repair transcription factor Sox9 was strongly enriched with GFP+ cells in the mutants compared with WT animals (Figure 5e, Supplementary Figure S9). Sox9+ cells were more likely to co-express phospho-histone H3 in the mutant animals, indicating a higher fraction of repairing cells successfully progressing through the cell cycle (Figure 5f, Supplementary Figure S9). These results indicate that loss of Pax2 and Pax8 protects against AKI early after injury, suggesting mutant proximal tubule cells acquire resistance to ischemia. Indeed, when we repeated these experiments in the very early stages of injury, 6 h after clamping, kidney function was better preserved in mutant mice (Figure 5g). This early protection also translated to a reduction in chronic injury. Both BUN and the expression of KIM-1 and Vcam1 were lower in the mutant mice 28 d after injury (Figure 5g and Supplementary Figure S10). These combined data suggest that improved chronic outcomes after IRI are mediated by inherent resistance to ischemic injury in the proximal-tubule-selective Pax2/8 double mutant mice.

**Figure 5.**
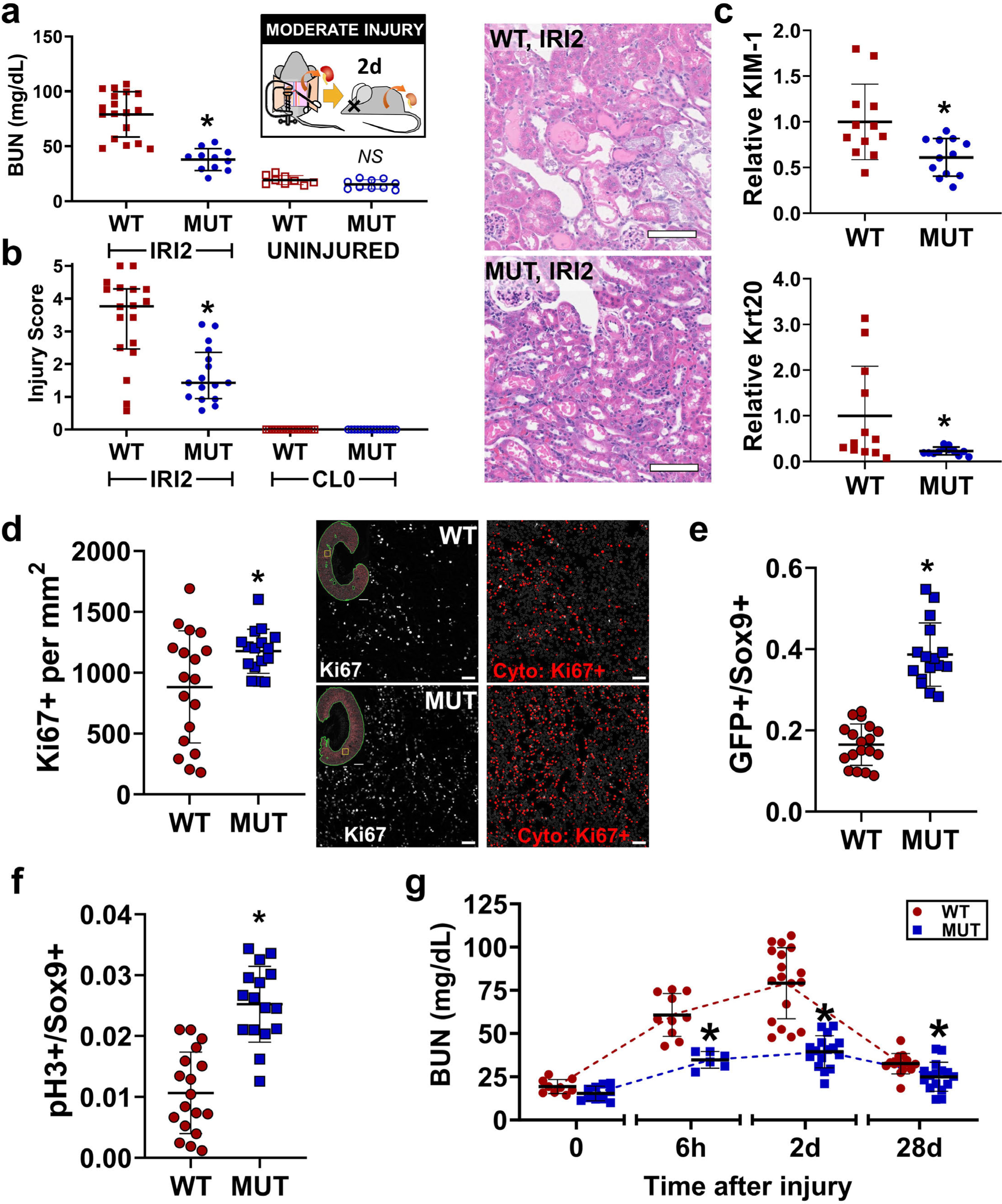
Pax2 and Pax8 mutant animals are protected from acute IRI. (**a**) BUN measured from terminal blood collection 2 d after unilateral IRI with contralateral nephrectomy (IRI2; *: *P* < 0.0001, Brown-Forsythe and Welch ANOVA for unequal variances with Dunnett’s post hoc test, *N =* 11-18 animals). (**b**) Acute injury score 2 d after IRI (IRI2) compared with uninjured kidneys collected at the time of surgery (CL0; *: *P* = 0.0005, Mann-Whitney test, IRI2 only, N-16-18, median ± interquartile range shown). Representative hematoxylin and eosin images are shown. Scale bar = 100 µm. (**c**) Relative expression of KIM-1 and Krt20 in whole kidney measured by real-time RT-PCR normalized to β-actin at 2 d after IRI (*: *P* = 0.01 and 0.03, respectively, t-test, *N =* 12). (**d**) Ki67+ cells measured using U-Net assisted cytometry over the cortex + OSOM 2 d after IRI (*: *P* = 0.02, t-test, *N =* 16-18). Representative images are shown in grayscale with corresponding cytometry results indicated in red (gray outlines represent negative nuclei). Scale bars: 1 mm (full section), 50 µm (inset). (**e**) Quantification of GFP+ cells in the subpopulation of Sox9+ cells analyzed over the cortex + OSOM (*: *P* < 0.0001, t-test, *N =* 16-18 animals). Representative Images are shown in Supplementary Figure S9. (**f**) Quantification of pH3+ cells in the Sox9+ subpopulation measured using the same methodology as in (**e**) (*: *P* < 0.0001, t-test, *N =* 16-18 animals). (**g**) BUN measured at various times after injury. Note time axis is not to scale (*: *P* = 0.0002, < 0.0001, and = 0.027 MUT vs WT for 6 h, 2 d, and 28 d respectively, Brown-Forsythe and Welch ANOVA for unequal variances with Dunnett’s post hoc test, *N =* 6-18). Strain abbreviations-WT: Pax2^wt/wt^, Pax8^wt/wt^, PEPCK Cre^+/y^, mT/mG^+/+^; MUT: Pax2^fl/fl^, Pax8^fl/fl^, PEPCK Cre^+/y^, mT/mG^+/+^.

### Pax2 and Pax8 mutant mice develop S3 proximal tubule cells with hallmarks of repair prior to injury

We next performed single nucleus RNA sequencing (snRNA seq) on uninjured WT and mutant kidneys to identify potential mechanisms and pathways underpinning the resistance to IRI conferred by Pax2 and Pax8 deletion (Supplementary Data S1). Contributions to each cluster were well-balanced between WT and mutant animals except for a new cluster that partially overlapped with S3 proximal tubule and was almost exclusively derived from mutant animals (arrow, Figure 6a, Supplementary Figure S11, Supplementary Figure S12). Furthermore, the number of differentially expressed genes between mutant and WT animals within each cluster was largest in the proximal tubule clusters, specifically the S3 segment (Supplementary Figure S12). There were few differentially expressed genes between WT and mutant mice in the endothelial, fibroblast, and immune clusters, suggesting that key differences explaining resistance to IRI were most likely cell autonomous in the proximal tubule. To explore this finding in detail, we extracted the subset of proximal tubule cells, including the mutant S3, and reclustered them into 5 distinct clusters (Figure 6b). We again obtained a new cluster near-exclusively derived from the mutant animals (Supplementary Figure S13).

**Figure 6.**
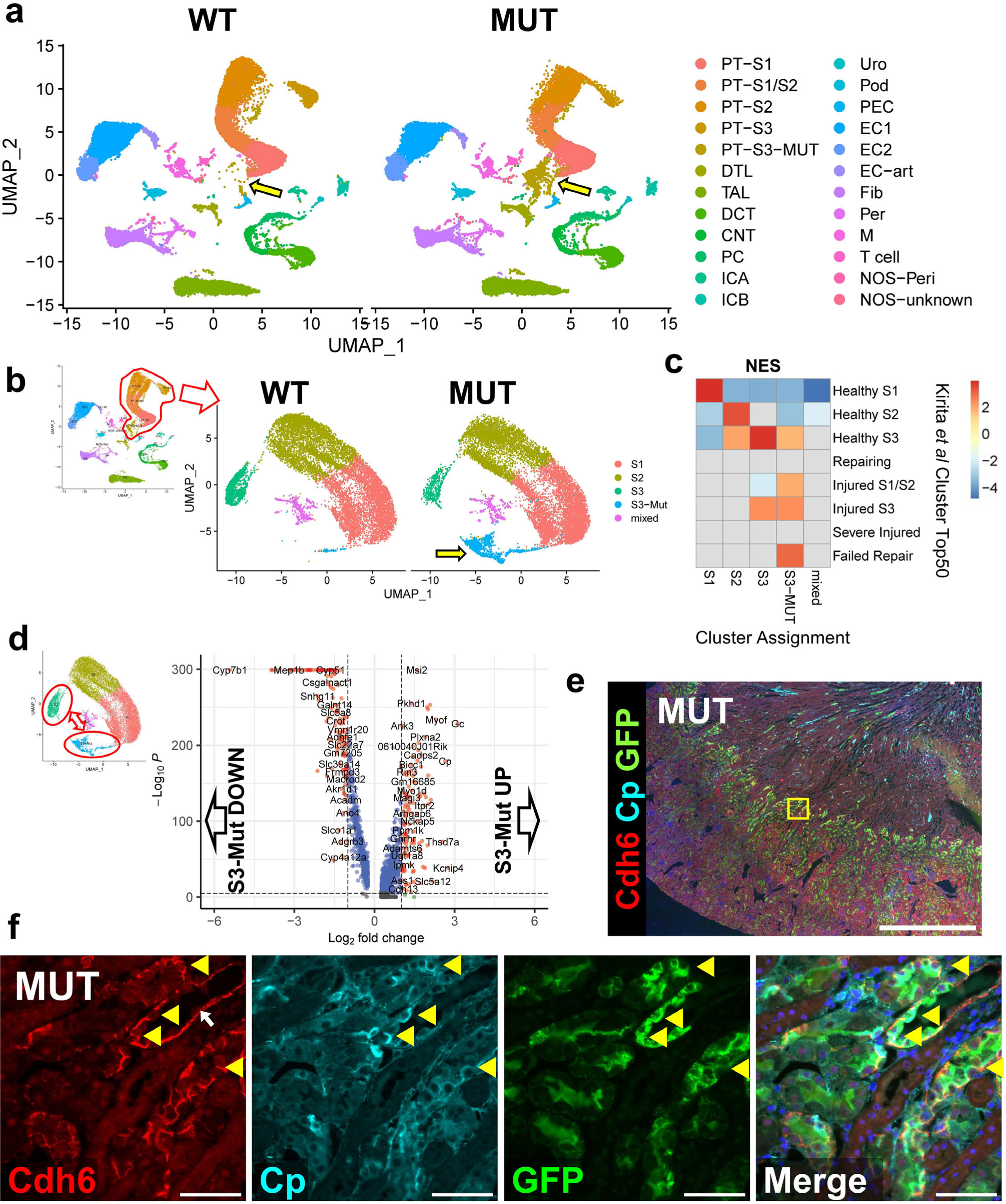
Single nucleus RNA sequencing identifies Pax2 and Pax8 mutant S3 proximal tubules with features of injury and repair in uninjured kidneys. (**a**) Uniform manifold approximation and projection (UMAP) plots of 57,670 high quality single nucleus transcriptomes from 4 WT and 4 mutant uninjured mice split by genotype. Cluster annotations derived as shown in Supplementary Figure S11. Arrow highlights the new populations of S3-like cells derived from the mutant mice. (**b**) Subclustering of only proximal tubule cells including the new mutant S3 cluster. Arrow highlights the cluster derived near-exclusively from mutant animals. (**c**) Gene set enrichment analysis (GSEA) of differentially expressed genes from clusters shown in (**b**) compared with the top 50 differentially expressed genes from each proximal tubule cluster in Kirita et al^6^. Data is presented as a heatmap of normalized enrichment scores (NES) for each comparison. Only enrichments with adjusted *P* < 0.05 are colored. Comparisons without significant enrichment are shown in gray. (**d**) Volcano plot of differentially expressed genes between mutant S3 (S3-Mut) and normal S3 (S3) clusters. (**e**) Wide field and (**f**) region of interest (ROI) show expression of cadherin 6 (Cdh6) at cell-cell junctions and membrane associated ceruloplasmin (Cp) in GFP+ cells, two genes that were differentially upregulated in S3-Mut cells and were localized to S3 proximal tubules. Location of the ROI shown is marked as a yellow box in (**e**). Yellow arrow heads indicate Cdh6+, Cp+, GFP+ cells. These cells were only observed in mutant animals. White arrow indicates cadherin 6 staining along the basement membrane, which was seen as the S3 transitions into the descending loop of Henle in both WT and MUT animals. Scale bars: 1 mm (wide field); 50 μm (ROI). Strain abbreviations-WT: Pax2^wt/wt^, Pax8^wt/wt^, PEPCK Cre^+/y^, mT/mG^+/+^; MUT: Pax2^fl/fl^, Pax8^fl/fl^, PEPCK Cre^+/y^, mT/mG^+/+^.

We hypothesized that genes involved in IRI repair may be differentially expressed in our mutant mice, explaining IRI resistance. Surprisingly, we found that the mutant-exclusive cluster strongly enriched in genes associated with chronic injury from multiple similar post-IRI datasets^6,13,28^ (Figure 6c and Supplementary Figure S13). These enriched signatures included a strong overlap with the population of epithelial cells found after AKI termed “failed repair” by Kirita et al^6^. However, in this context the “failed repair” signature did not appear to compromise renal function and thus may be more related to a preconditioned phenotype. To corroborate these findings, we performed bulk RNA sequencing of uninjured proximal tubules purified by magnetic activated cell sorting (MACS) with lotus lectin (Supplementary Figure S14, Supplementary Data S2). Likewise, mutant proximal tubules isolated with this technique were enriched with the “failed repair” signature and depleted of healthy S2 and S3 proximal tubule genes (Supplementary Figure S14). Notably, Vcam1, widely used to mark “failed repair” cells^6,13^, was not upregulated in the mutant proximal tubule in either analysis.

To further characterize this cell population, we used PROGENy^29–31^ to infer the activity of well-characterized signaling pathways. This approach revealed increased activity of NFkB, TNFα, JAK-STAT, EGFR, and p53 signaling in the mutant S3 population and decreased activity of androgen signaling (Supplementary Figure S15). Likewise, we found increased activity of the transcription factors Smad1, Klf6, and Relb in mutant S3 and decreased activity of Rxra using signatures defined by DoRothEA^32^ (Supplementary Figure S16). These pathways are associated with injury and recovery after ischemic AKI^6,33,34^. However, pseudotime trajectory analysis using Slingshot^35^ did not reveal a direct path from normal S3 to the mutant S3 population (Supplementary Figure S17). This finding suggested that Pax2 and Pax8 loss induced cell-autonomous reprograming rather than a response to injury where a continuum of cell states has been observed^6,13^. Furthermore, inflammatory markers that are upregulated after injury, such as Ccl2, Cxcl1, and Cxcl2^6,13^, were not upregulated in the mutant S3 cells.

Based on these findings, we focused our subsequent analysis on key differences between normal S3 and mutant S3 cells specifically. Among 269 differentially expressed genes (Figure 6d), we selected Cp (ceruloplasmin), Gc (vitamin D binding protein), Cdh6 (Cadherin 6), and Aldh1a1 (aldehyde dehydrogenase 1-A1) as candidate marker genes, due to their relative specificity for the mutant S3 cluster (Supplementary Figure S18) and concomitant upregulation in our MACS-purified proximal tubule RNA-seq dataset (Supplementary Figure S14). In uninjured mutant animals, but not WT controls, we specifically identified a population of cells only in the S3 segment with cadherin 6 expression, membrane bound ceruloplasmin, and universal expression of GFP (Figure 6e-f). Similarly, cells strongly expressing Aldh1a1 and Gc were identified in S3 proximal tubule of mutant mice, though we found Aldh1a1 protein was also expressed in other proximal tubule segments exclusively in mutants (Supplementary Figure S19). These results indicate that the S3-Mut population identified by snRNA seq localizes to S3 proximal tubule and is derived from GFP+ Pax2 and Pax8 mutant cells. Interestingly, we also observed that the repair markers Ki67 and Sox9 were markedly increased in the GFP+ proximal tubule populations of uninjured mutant animals, predominantly in the S3 segment (Figure 7). Many of these features are shared with immature proximal tubules in development, such as Cdh6 expression^13,36^, suggesting that Pax2 and Pax8 depletion may induce an immature progenitor-like state that could explain resistance to injury. However, we found no significant positive enrichment of early developing proximal tubule gene signatures^13,37,38^ in the mutant S3 population (Supplementary Figure S20). Together these data show that mutant mice develop a unique population of S3 proximal tubule cells that express genes associated with ischemic injury and activate repair-associated processes even before injury. These findings seem to suggest an adaptive role for many genes associated with the “failed repair” phenotype and is reminiscent of other states where pre-injury stress responses protect against ischemic AKI.

**Figure 7.**
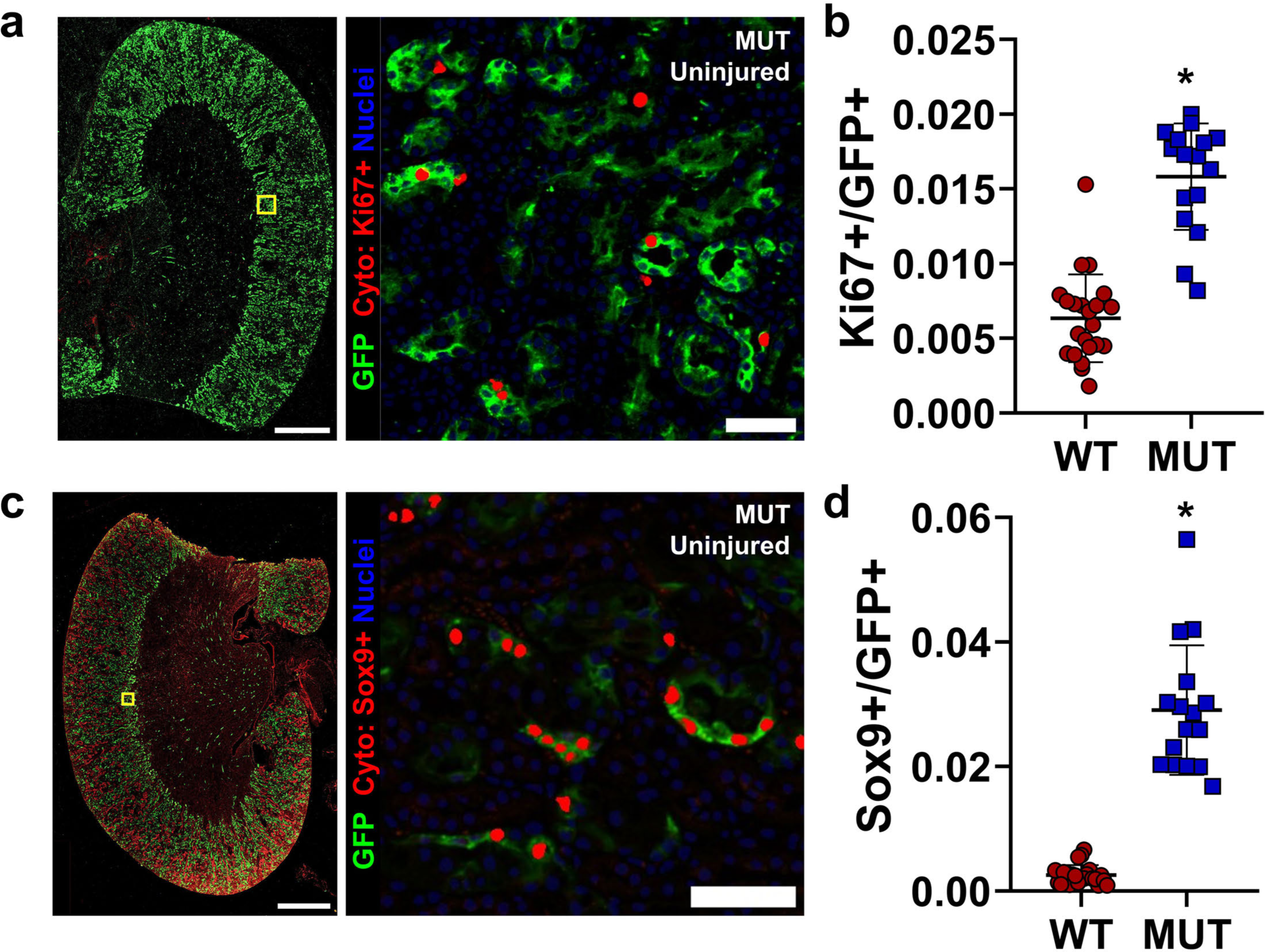
Mutant Pax2 and Pax8 proximal tubules show increased turnover and repair in uninjured kidneys. (**a**) Representative co-stained images for Ki67 and GFP in uninjured kidneys and a representative region of interest (ROI) near the S3 segment showing GFP staining and Ki67 nuclei as determined by deep-learning-assisted tissue cytometry (Cyto, in red). Note one pole of the kidney was intentionally removed for other analyses. (**b**) Quantification of the fraction of GFP+ cells that co-stained with Ki67 over the cortex + OSOM (*: *P* < 0.0001, t-test, *N =* 16-22 animals). (**c**) Representative staining for Sox9 as in (**a**). Note nonspecific, non-nuclear staining for Sox9 in the outer cortex was ignored by deep-learning-assisted cytometry. (**d**) Quantification of the fraction of GFP+ cells that co-stained for nuclear Sox9 detected over the cortex + OSOM (*: *P* < 0.0001, t-test, *N =* 16-22 animals). Scale bars: 1 mm (wide field); 50 μm (ROI). Strain abbreviations-WT: Pax2^wt/wt^, Pax8^wt/wt^, PEPCK Cre^+/y^, mT/mG^+/+^; MUT: Pax2^fl/fl^, Pax8^fl/fl^, PEPCK Cre^+/y^, mT/mG^+/+^.

### Pax2 and Pax8 mutant S3 proximal tubule cells activate conserved mechanisms of resistance to ischemic injury

To define potential mechanisms underpinning resistance to IRI, we compared the expression profile of mutant S3 cells with pre-injury expression data from hypoxic preconditioning and caloric restriction^39^, which also confer protection against IRI. We noted a profound enrichment in genes associated with both interventions in the mutant S3 cells (Figure 8a-b). Female sex also confers resistance to IRI in mice^40^. We found similar strong enrichment in genes expressed in female proximal tubules^41^ in our mutant S3 (Figure 8c-d). Between these diverse data sets, we identified 12 genes upregulated and 24 downregulated in mutant S3 cells and at least two of the other data sets. We noted an increase in genes associated with oxidative stress among the differentially upregulated genes (e.g. Gsta2, Gsta3, Cp, Kynu; Supplementary Figure S21). These results indicate that the loss of Pax2 and Pax8 in proximal tubule cells induces conserved mechanisms of resistance to IRI and suggests that confining these changes to the S3 segment of the proximal tubule may be sufficient to mitigate ischemic AKI.

**Figure 8.**
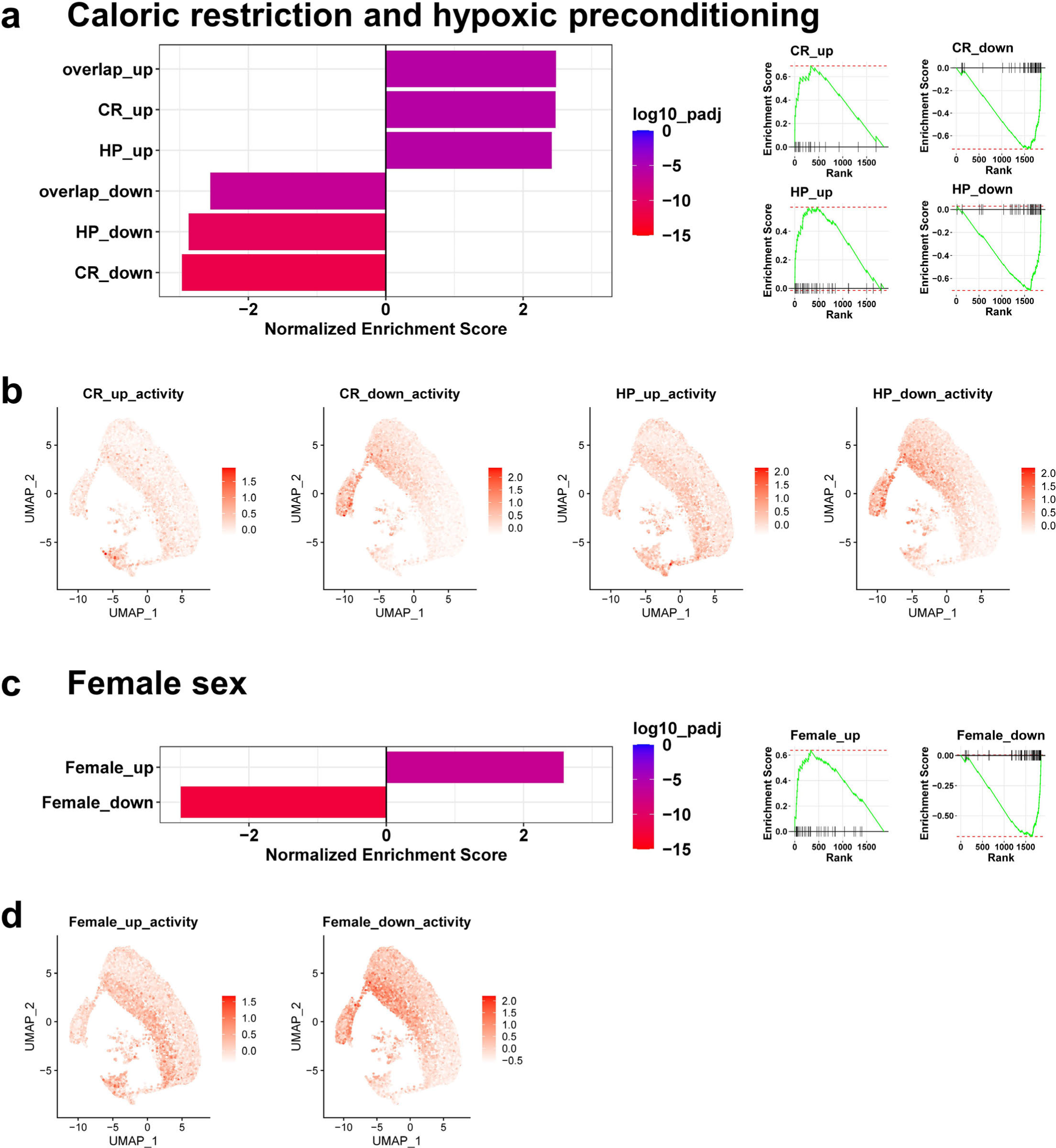
Mutant S3 proximal tubule cells activate conserved programs of resistance to ischemic AKI. (**a**) Bar plots of normalized enrichment scores for the top 100 differentially up- or down-regulated genes induced by caloric restriction (CR), hypoxic preconditioning (HP), or both (overlap) obtained from whole kidney RNA seq^39^ in the genes differentially expressed in uninjured mutant S3 proximal tubule relative to healthy S3 in our studies (shown in Figure 6d). GSEA enrichment plots are shown right. (**b**) Activity of the indicated gene set (as determined by the average expression z-score of set genes) for individual cells mapped to the UMAP shown in Figure 6. (**c**) Normalized enrichment scores and GSEA enrichment plots as in (**a**) comparing the top 100 up- or down-regulated genes in female proximal tubule cells compared with males^41^ with the differentially expressed genes in mutant S3 proximal tubule in our studies. (**d**) Cell-wise gene set activity as in (**b**) for female sex gene sets.

## DISCUSSION

In this report we examined the role of Pax2 and Pax8 in repair after ischemic AKI. Surprisingly, proximal tubule conditional Pax2/8 double mutant mice showed enhanced regeneration and recovery after IRI, mediated by a resistance to injury. Prior to injury, we found mutant mice develop a unique population of S3 proximal tubule cells with transcriptional signatures that overlap with injury response and with multiple mechanisms of protection against ischemic AKI. These findings provide new perspectives on the roles of Pax proteins in regeneration and their functions in cells that display evidence of incomplete repair, two areas of central importance to the development of therapies for ischemic AKI.

Pax proteins, most commonly Pax2, have long been used as markers associated with regeneration in the kidney^4,11,42,43^. Until recently, Pax8 was not thought to be involved in AKI recovery^43,44^, though recent studies with newer techniques now suggest Pax8 is dynamically regulated during repair^6,12,13,28,33^. In addition, human^45,46^ and rodent^42^ proximal tubules contains scattered cells marked by higher levels of Pax2 and/or Pax8 that may have progenitor-like properties and may contribute disproportionately to repair. Work in the early 2000’s suggested that Pax2 was involved in maintaining proliferation and preventing apoptosis of injured cells^43,47,48^. The transient upregulation of Pax2 and Pax8 in normal repair^11,43^ and the developmental role of Pax2 and Pax8 mediating mesenchymal-to-epithelial transitions^49^ are consistent with this role in AKI recovery. Furthermore, we recently showed that Paxip1, which encodes the Pax transactivation-domain interacting protein (PTIP), is required for successful repair after AKI^25^ with critical implications for treatment of ischemic AKI. Recently, Sako et al reported impaired recovery of Pax2 conditional knockout mice driven by kidney androgen-regulated protein (Kap) Cre, attributed to decreased proliferation and increased apoptosis of mutant cells^15^. These results implicated Pax2 and Pax8 as critical survival factors after injury^50^ Based on these observations, we initially anticipated that Pax2/8 double mutants would show severely impaired AKI recovery.

However, in this report we showed that Pax2 and Pax8 expression was associated with incomplete repair phenotypes and that Pax2/8 loss protects against chronic injury after IRI. Our initial explanation for this finding was that severely injured Pax mutant cells die or fail to proliferate, whereas severely injured cells that retain Pax function survive albeit in a chronically injured state. However, in early injury we found increased evidence of proliferation and repair in mutant cells, and we noted differences in function even 6 h after IRI, suggesting the dominant effect is a resistance to injury. Even before injury, Pax2/8 deletions lead to extensive cell autonomous reprogramming specifically in the S3 proximal tubule, increased turnover, and features of repair. These findings highlight a new unappreciated role for Pax proteins in the uninjured proximal tubule and suggest that, while Pax2 and Pax8 may mark progenitor-like cells in the nephron, they are dispensable for normal repair.

Our findings contrast with those of Sako et al^15^. This discrepancy could be attributable to dual deletion of Pax2 and Pax8 or use of the PEPCK Cre driver in our studies. We noted that Kap, used to drive Cre in their report, was among the small subset of genes downregulated in our Pax2 and Pax8 mutant S3 cells that also overlapped with expression changes in caloric restriction and female proximal tubule. Thus, it is a possible target of Pax protein regulation in mediating sensitivity to AKI. This coincidental and likely unappreciated feature makes Kap-Cre poorly suited for studying Pax protein function in ischemic AKI. In contrast, Pck1 which encodes PEPCK was not differentially expressed between mutant and WT S3 proximal tubules.

Genes associated with damage and incomplete repair are correlated with fibrosis, suggesting that this signature may be deleterious. Surprisingly, we found these signatures were upregulated in Pax2/8 mutant S3 proximal tubule cells prior to injury and, in this context, protect against ischemic injury and fibrosis. Further characterization of the transcriptional signature of these cells revealed strong overlapping features with signatures derived from mouse models of caloric restriction or hypoxia before the induction of AKI. Female mice, which are more resistant to AKI, also exhibit similar signatures^39^. These findings suggest common features of AKI resistance that may be attenuated, at least in part, by Pax2 and Pax8. While many of these altered genes have an obvious connection to ischemic stress (e.g. Gsta2 and Gsta3), many do not. In fact, one such gene, Kynu, was recently identified via an alternative approach studying hypoxic preconditioning and mediates its effect by preserving NAD+^51^. Our overlapping gene sets also include the transporters Slc7a12 and Slc6A6 which are transiently upregulated during recovery from ischemic injury, implying an adaptive function^33^. However, it is notable that neither our mutant S3 nor other resistance gene sets showed increased expression of Vcam1, which has been strongly associated with maladaptive repair^13,14^, suggesting injured cells can activate both adaptive and maladaptive transcriptional programs and that Pax2/8 may regulate the balance.

In this study, we wanted to limit Pax deletions to the proximal tubule, a key site of regeneration after AKI. Thus, we utilized the PEPCK-Cre driver strain because it activates late in nephrogenesis in proximal tubule segments, so that Pax2/8 double mutants do not exhibit the significant renal pathology or loss of function observed in global mutants^21^. Since the PEPCK-Cre driver is constitutive, several caveats must be considered. Pax2/8 mutant cells may be more sensitive to cell death. The increased baseline proliferation in uninjured double mutants could be due to replacement of mutant cells by cells that have not deleted Pax2/8 because they do not yet express PEPCK. As cells then replace mutants, they activate PEPCK and delete the Pax genes. Thus, this low-level stress and replacement may underlie the increased resistance to AKI. This also could explain the residual level of Pax protein expression we see in focal areas.

In summary, the data presented are contrary to what we might have predicted based on the need for Pax proteins in developing nephrons and the enhanced expression of Pax proteins during regeneration, which is thought to mimic aspects of development. Rather, deletion of Pax2 and Pax8 in mature S3 proximal tubule cells did not significantly impact renal function but did alter their transcriptional signature to mimic a stress-induced phenotype, similar to that seen after global ischemic preconditioning or caloric restriction. This stress-induced signature increased resistance to primary ischemic damage and reduced the number of cells going on to develop the failed repair phenotype that characterizes the AKI-to-CKD transition. Better understanding of this stress induced signature and ways of inducing it selectively could improve strategies for mitigating acute and chronic renal injury.

## DISCLOSURE

All authors declare no competing interests.

## ACKNOWLEGEMENTS

Support for this work was provided by the University of Michigan O’Brien Kidney Center (DK-P30-081943 to J.A.B.), NIH K08 DK125776 (to J.A.B.), NIH R01 DK054740 to (G.R.D.), NIH R01DK073722 to (G.R.D.), and a generous gift from Josh and Audrey Rumsey in memory of Isaiah. We would also like to acknowledge the support from the Bioinformatics Core (RRID:SCR_019168) and Tissue and Molecular Pathology Core at the University of Michigan School of Medicine. We are grateful to Phillip McCown for assistance with coding associated with snRNAseq data analysis.

## AUTHOR CONTRIBUTIONS

JAB and GRD designed the research. JAB, ACT, MCM, AA-S, MC, JFS, EAO, and AS performed experiments. JAB and RM analyzed snRNA sequencing data. JAB, JMW, SRP, and GRD analyzed and interpreted results. JAB wrote the manuscript with contributions from all authors.

## DATA SHARING STATEMENT

RNA sequencing data was deposited on Mendeley Data (doi:10.17632/jdcbrjx3pp.1). Algorithms for image analysis are available at https://github.com/Beamish-Lab/High-throughput-Image-Analysis-of-Kidney-Injury.

## Notes

### Competing Interest Statement

The authors have declared no competing interest.

## REFERENCES

1. Chertow GM, Burdick E, Honour M, Bonventre JV, Bates DW. Acute Kidney Injury, Mortality, Length of Stay, and Costs in Hospitalized Patients. Journal of the American Society of Nephrology. 2005;16(11):3365–3370. doi:10.1681/ASN.2004090740

2. Coca SG, Yusuf B, Shlipak MG, Garg AX, Parikh CR. Long-term Risk of Mortality and Other Adverse Outcomes After Acute Kidney Injury: A Systematic Review and Meta-analysis. American Journal of Kidney Diseases. 2009;53(6):961–973. doi:10.1053/j.ajkd.2008.11.034 PMID - 19346042

3. Bonventre JV, Yang L. Cellular pathophysiology of ischemic acute kidney injury. Journal of Clinical Investigation. 2011;121(11):4210–4221. doi:10.1172/JCI45161

4. Kusaba T, Lalli M, Kramann R, Kobayashi A, Humphreys BD. Differentiated kidney epithelial cells repair injured proximal tubule. Proceedings of the National Academy of Sciences. 2014;111(4):1527–1532. doi:10.1073/pnas.1310653110

5. Ferenbach DA, Bonventre JV. Mechanisms of maladaptive repair after AKI leading to accelerated kidney ageing and CKD. Nature Reviews Nephrology. 2015;11(5):264–276. doi:10.1038/nrneph.2015.3

6. Kirita Y, Wu H, Uchimura K, Wilson PC, Humphreys BD. Cell profiling of mouse acute kidney injury reveals conserved cellular responses to injury. Proceedings of the National Academy of Sciences. 2020;117(27):15874–15883. doi:10.1073/pnas.2005477117 PMID - 32571916

7. Yang L, Besschetnova TY, Brooks CR, Shah JV, Bonventre JV. Epithelial cell cycle arrest in G2/M mediates kidney fibrosis after injury. Nature Medicine. 2010;16(5):535–543. doi:10.1038/nm.2144 PMID - 20436483

8. Venkatachalam MA, Weinberg JM, Kriz W, Bidani AK. Failed Tubule Recovery, AKI-CKD Transition, and Kidney Disease Progression. Journal of the American Society of Nephrology. 2015;26(8):1765–1776. doi:10.1681/ASN.2015010006

9. Chiba T, Hukriede N, de Caestecker MP. Kidney Regeneration: Lessons from Development. Current Pathobiology Reports. 2015;3(1):67–79. doi:10.1007/s40139-015-0069-z

10. Little M, Kairath P. Does Renal Repair Recapitulate Kidney Development? Journal of the American Society of Nephrology. 2016;28(1):34–46. doi:10.1681/ASN.2016070748

11. Imgrund M, Gröne E, Gröne HJ, et al. Re-expression of the developmental gene Pax-2 during experimental acute tubular necrosis in mice1. Kidney International. 1999;56(4):1423–1431-1431. doi:10.1046/j.1523-1755.1999.00663.x

12. Kang HM, Huang S, Reidy K, Han SH, Chinga F, Susztak K. Sox9-Positive Progenitor Cells Play a Key Role in Renal Tubule Epithelial Regeneration in Mice. Cell Reports. 2016;14(4):861–871-871. doi:10.1016/j.celrep.2015.12.071

13. Ide S, Kobayashi Y, Ide K, et al. Ferroptotic stress promotes the accumulation of pro-inflammatory proximal tubular cells in maladaptive renal repair. eLife. 2021;10:e68603. doi:10.7554/elife.68603 PMID - 34279220

14. Gerhardt LMS, Koppitch K, Gestel Jv, et al. Lineage Tracing and Single-Nucleus Multiomics Reveal Novel Features of Adaptive and Maladaptive Repair after Acute Kidney Injury. Journal of the American Society of Nephrology. 2023;Publish Ahead of Printdoi:10.1681/asn.0000000000000057 PMID - 36735940

15. Sako K, Furuichi K, Makiishi S, et al. Cyclin-dependent kinase 4-related tubular epithelial cell proliferation is regulated by Paired box gene 2 in kidney ischemia-reperfusion injury. Kidney International. 2022;102(1):45–57. doi:10.1016/j.kint.2022.03.022 PMID - 35483529

16. Grimley E, Dressler GR. Are Pax proteins potential therapeutic targets in kidney disease and cancer? Kidney International. 2018;(Am J Kidney Dis 42 2003)doi:10.1016/j.kint.2018.01.025

17. Patel SR, Kim D, Levitan I, Dressler GR. The BRCT-domain containing protein PTIP links PAX2 to a histone H3, lysine 4 methyltransferase complex. Dev Cell. Oct 2007;13(4):580–92. doi:10.1016/j.devcel.2007.09.004

18. Abraham S, Paknikar R, Bhumbra S, et al. The Groucho-associated Phosphatase PPM1B Displaces Pax Transactivation Domain Interacting Protein (PTIP) to Switch the Transcription Factor Pax2 from a Transcriptional Activator to a Repressor. Journal of Biological Chemistry. 2015;290(11):7185–7194-7194. doi:10.1074/jbc.m114.607424

19. Soofi A, Levitan I, Dressler GR. Two novel EGFP insertion alleles reveal unique aspects of Pax2 function in embryonic and adult kidneys. Developmental Biology. 2012;365(1):241–250-250. doi:10.1016/j.ydbio.2012.02.032

20. Bouchard M, Souabni A, Mandler M, Neubüser A, Busslinger M. Nephric lineage specification by Pax2 and Pax8. Genes Dev. Nov 2002;16(22):2958–70. doi:10.1101/gad.240102

21. Laszczyk AM, Higashi AY, Patel SR, et al. Pax2 and Pax8 Proteins Regulate Urea Transporters and Aquaporins to Control Urine Concentration in the Adult Kidney. Journal of the American Society of Nephrology. 2020;31(6):1212–1225. doi:10.1681/asn.2019090962 PMID - 32381599

22. Mehta RL, Pascual MT, Soroko S, et al. Spectrum of acute renal failure in the intensive care unit: The PICARD experience. Kidney International. 2004;66(4):1613–1621. doi:10.1111/j.1523-1755.2004.00927.x PMID - 15458458

23. Liaño F, Pascual J, Group TMARFS. Epidemiology of acute renal failure: A prospective, multicenter, community-based study. Kidney International. 1996;50(3):811–818. doi:10.1038/ki.1996.380 PMID - 8872955

24. Rankin EB, Tomaszewski JE, Haase VH. Renal Cyst Development in Mice with Conditional Inactivation of the von Hippel-Lindau Tumor Suppressor. Cancer Research. 2006;66(5):2576–2583-2583. doi:10.1158/0008-5472.can-05-3241

25. Soofi A, Kutschat AP, Azam MH, Laszczyk AM, Dressler GR. Regeneration after acute kidney injury requires PTIP mediated epigenetic modifications. JCI Insight. 2020;5(3)doi:10.1172/jci.insight.130204 PMID - 31917689

26. Hao Y, Hao S, Andersen-Nissen E, et al. Integrated analysis of multimodal single-cell data. Cell. 2021;184(13):3573–3587.e29. doi:10.1016/j.cell.2021.04.048 PMID - 34062119

27. McElliott MC, Al-Suraimi A, Telang AC, et al. High-throughput image analysis with deep learning captures heterogeneity and spatial relationships after kidney injury. Scientific Reports. 2023;13(1):6361. doi:10.1038/s41598-023-33433-3 PMID - 37076596

28. Gerhardt LMS, Liu J, Koppitch K, Cippà PE, McMahon AP. Single-nuclear transcriptomics reveals diversity of proximal tubule cell states in a dynamic response to acute kidney injury. Proceedings of the National Academy of Sciences of the United States of America. 2021;118(27):e2026684118. doi:10.1073/pnas.2026684118 PMID - 34183416

29. Holland CH, Szalai B, Saez-Rodriguez J. Transfer of regulatory knowledge from human to mouse for functional genomics analysis. Biochimica et Biophysica Acta (BBA) - Gene Regulatory Mechanisms. 2020;1863(6):194431. doi:10.1016/j.bbagrm.2019.194431 PMID - 31525460

30. Holland CH, Tanevski J, Perales-Patón J, et al. Robustness and applicability of transcription factor and pathway analysis tools on single-cell RNA-seq data. Genome Biology. 2020;21(1):36. doi:10.1186/s13059-020-1949-z PMID - 32051003

31. Schubert M, Klinger B, Klünemann M, et al. Perturbation-response genes reveal signaling footprints in cancer gene expression. Nature Communications. 2018;9(1):20. doi:10.1038/s41467-017-02391-6 PMID - 29295995

32. Garcia-Alonso L, Holland CH, Ibrahim MM, Turei D, Saez-Rodriguez J. Benchmark and integration of resources for the estimation of human transcription factor activities. Genome Research. 2019;29(8):1363–1375. doi:10.1101/gr.240663.118 PMID - 31340985

33. Li H, Dixon EE, Wu H, Humphreys BD. Comprehensive single-cell transcriptional profiling defines shared and unique epithelial injury responses during kidney fibrosis. Cell Metabolism. 2022;34(12):1977–1998.e9. doi:10.1016/j.cmet.2022.09.026 PMID - 36265491

34. Piret SE, Guo Y, Attallah AA, et al. Krüppel-like factor 6–mediated loss of BCAA catabolism contributes to kidney injury in mice and humans. Proceedings of the National Academy of Sciences. 2021;118(23):e2024414118. doi:10.1073/pnas.2024414118 PMID - 34074766

35. Street K, Risso D, Fletcher RB, et al. Slingshot: cell lineage and pseudotime inference for single-cell transcriptomics. BMC Genomics. 2018;19(1):477. doi:10.1186/s12864-018-4772-0

36. Marable SS, Chung E, Park J-S. Hnf4a Is Required for the Development of Cdh6-Expressing Progenitors into Proximal Tubules in the Mouse Kidney. Journal of the American Society of Nephrology. 2020;31(11):2543–2558. doi:10.1681/asn.2020020184 PMID - 32764140

37. Adam M, Potter AS, Potter SS. Psychrophilic proteases dramatically reduce single-cell RNA-seq artifacts: a molecular atlas of kidney development. Development. 10 2017;144(19):3625–3632. doi:10.1242/dev.151142

38. Combes AN, Phipson B, Lawlor KT, et al. Single cell analysis of the developing mouse kidney provides deeper insight into marker gene expression and ligand-receptor crosstalk. Development. 2019;146(12):dev178673. doi:10.1242/dev.178673 PMID - 31118232

39. Johnsen M, Kubacki T, Yeroslaviz A, et al. The Integrated RNA Landscape of Renal Preconditioning against Ischemia-Reperfusion Injury. Journal of the American Society of Nephrology. 2020;31(4):716–730. doi:10.1681/asn.2019050534 PMID - 32111728

40. Park KM, Kim JI, Ahn Y, Bonventre AJ, Bonventre JV. Testosterone Is Responsible for Enhanced Susceptibility of Males to Ischemic Renal Injury*. Journal of Biological Chemistry. 2004;279(50):52282–52292. doi:10.1074/jbc.m407629200 PMID - 15358759

41. Ide S, Ide K, Abe K, et al. Sex differences in resilience to ferroptosis underlie sexual dimorphism in kidney injury and repair. Cell Reports. 2022;41(6):111610. doi:10.1016/j.celrep.2022.111610 PMID - 36351395

42. Lazzeri E, Angelotti ML, Peired A, et al. Endocycle-related tubular cell hypertrophy and progenitor proliferation recover renal function after acute kidney injury. Nature Communications. 2018;9(1):1344. doi:10.1038/s41467-018-03753-4

43. Maeshima A, Maeshima K, Nojima Y, Kojima I. Involvement of Pax-2 in the action of activin A on tubular cell regeneration. J Am Soc Nephrol. Dec 2002;13(12):2850–9. doi:10.1097/01.asn.0000035086.93977.e9

44. Lindoso RS, Verdoorn KS, Einicker-Lamas M. Renal recovery after injury: the role of Pax-2. Nephrology Dialysis Transplantation. 2009;24(9):2628–2633. doi:10.1093/ndt/gfp307 PMID - 19556301

45. Smeets B, Boor P, Dijkman H, et al. Proximal tubular cells contain a phenotypically distinct, scattered cell population involved in tubular regeneration. The Journal of Pathology. 2013;229(5):645–659. doi:10.1002/path.4125 PMID - 23124355

46. Eymael J, Broek Mvd, Miesen L, et al. Human scattered tubular cells represent a heterogeneous population of glycolytic dedifferentiated proximal tubule cells. The Journal of Pathology. 2022;doi:10.1002/path.6029 PMID - 36373978

47. Zhang SL, Guo J, Moini B, Ingelfinger JR. Angiotensin II stimulates Pax-2 in rat kidney proximal tubular cells: impact on proliferation and apoptosis. Kidney Int. Dec 2004;66(6):2181–92. doi:10.1111/j.1523-1755.2004.66008.x

48. Torban E, Eccles MR, Favor J, Goodyer PR. PAX2 suppresses apoptosis in renal collecting duct cells. Am J Pathol. Sep 2000;157(3):833–42. doi:10.1016/S0002-9440(10)64597-X

49. Rothenpieler UW, Dressler GR. Pax-2 is required for mesenchyme-to-epithelium conversion during kidney development. *Development (Cambridge*, England). 1993;119(3):711–20.

50. Cohen T, Loutochin O, Amin M, Capolicchio J-P, Goodyer P, Jednak R. PAX2 is reactivated in urinary tract obstruction and partially protects collecting duct cells from programmed cell death. American Journal of Physiology-Renal Physiology. 2007;292(4):F1267–F1273. doi:10.1152/ajprenal.00281.2006 PMID - 17164400

51. Torosyan R, Huang S, Bommi PV, et al. Hypoxic preconditioning protects against ischemic kidney injury through the IDO1/kynurenine pathway. Cell Reports. 2021;36(7):109547. doi:10.1016/j.celrep.2021.109547 PMID - 34407414

52. Falk T, Mai D, Bensch R, et al. U-Net: deep learning for cell counting, detection, and morphometry. Nature Methods. 2019;16(1):67–70. doi:10.1038/s41592-018-0261-2 PMID - 30559429

53. Badia-i-Mompel P, Santiago JV, Braunger J, et al. decoupleR: ensemble of computational methods to infer biological activities from omics data. Bioinformatics Advances. 2022;2(1):vbac016. doi:10.1093/bioadv/vbac016 PMID - 36699385

54. Marotta P, Amendola E, Scarfò M, et al. The paired box transcription factor Pax8 is essential for function and survival of adult thyroid cells. Molecular and Cellular Endocrinology. 2014;396(1-2):26–36-36. doi:10.1016/j.mce.2014.08.004

55. Borza CM, Bolas G, Bock F, et al. DDR1 contributes to kidney inflammation and fibrosis by promoting the phosphorylation of BCR and STAT3. JCI Insight. 2021;doi:10.1172/jci.insight.150887 PMID - 34941574

56. Chen J, Chen J-K, Conway EM, Harris RC. Survivin Mediates Renal Proximal Tubule Recovery from AKI. Journal of the American Society of Nephrology. 2013;24(12):2023–2033. doi:10.1681/asn.2013010076 PMID - 23949800

57. McGinnis CS, Murrow LM, Gartner ZJ. DoubletFinder: Doublet Detection in Single-Cell RNA Sequencing Data Using Artificial Nearest Neighbors. Cell Systems. 2019;8(4):329–337.e4. doi:10.1016/j.cels.2019.03.003 PMID - 30954475

58. Korotkevich G, Sukhov V, Budin N, Shpak B, Artyomov MN, Sergushichev A. Fast gene set enrichment analysis. bioRxiv. 2021:060012. doi:10.1101/060012

59. Love MI, Huber W, Anders S. Moderated estimation of fold change and dispersion for RNA-seq data with DESeq2. Genome Biology. 2014;15(12):550. doi:10.1186/s13059-014-0550-8 PMID - 25516281

