## Supplementary for "Pax Protein Depletion in Proximal Tubules Triggers Conserved Mechanisms of Resistance to Acute Ischemic Kidney Injury and Prevents Transition to Chronic Kidney Disease"

**TABLE OF CONTENTS:**

**Supplementary Figure S1. Detailed overview of mouse strain genetics.**

**Supplementary Figure S2. Overview of deep-learning-assisted tissue cytometry to measure proximal tubule cell Pax protein expression after injury**.

**Supplementary Figure S3. Correlation of Pax2 or Pax8 with KIM-1 expression in GFP+ proximal tubule cells**.

**Supplementary Figure S4. Pax2 and Pax8 are depleted in GFP expressing cells of mutant mice.**

**Supplementary Figure S5. Both Pax2 and Pax8 are depleted in S3 proximal tubule.**

**Supplementary Figure S6. Mutant animals show normal gross histology and similar kidney function compared with WT controls**.

**Supplementary Figure S7. Pax2 and Pax8 mutant cells are less likely to contribute to incomplete repair marked by KIM-1**.

**Supplementary Figure S8. Pax2 and Pax8 mutant mice with alternative genetics are also protected from chronic injury after severe IRI.**

**Supplementary Figure S9. Supporting data for Figure 5.**

**Supplementary Figure S10. Pax2 and Pax8 mutant mice develop less chronic injury after IRI.**

**Supplementary Figure S11. Cluster alignment of whole kidney snRNA seq.**

**Supplementary Figure S12. Cluster composition and differentially expressed genes from whole kidney snRNA seq.**

**Supplementary Figure S13. Composition of subclustered proximal tubule cells and gene set enrichment (GSEA) with alternative single cell/single nucleus RNA sequencing data sets.**

**Supplementary Figure S14. Purification and transcriptional profiling of proximal tubules using magnetic activated cell sorting (MACS)**.

**Supplementary Figure S15. Proximal tubule pathway activation using PROGENy**.

**Supplementary Figure S16. Transcription factor activity in proximal tubule clusters using DoRothEA**.

**Supplementary Figure S17. Pseudotime trajectory analysis of proximal tubule cells.**

**Supplementary Figure S18. Expression of selected markers of mutant S3 proximal tubule by scRNA seq.**

**Supplementary Figure S19. Expression of alternative markers of mutant S3 proximal tubule by immunofluorescence.**

**Supplementary Figure S20. Mutant S3 cells do not show positive enrichment of developmental markers.**

**Supplementary Figure S21. Shared genes altered in Pax2 and Pax8 mutant S3 proximal tubule also differentially expressed in other mechanisms of IRI protection.**

**Supplementary Figure S22. BUN correlates with histologic injury and serum creatinine.**

**Supplementary Methods.**

**Supplementary Table S1. Genotyping PCR Primers**

**Supplementary Table S2. Antibodies**

**Supplementary Table S3. Quantitative PCR Primers**

**Supplementary Data S1. Parse_snRNAseq_data.xlsx**

**Supplementary Data S2. Purified_PT_RNAseq_data.xlsx**

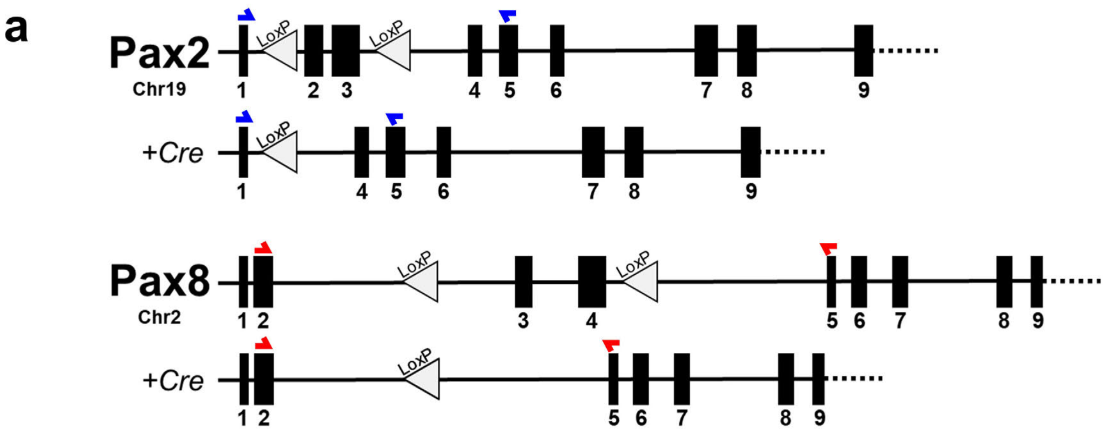


b

| Strain Abbreviation | Pax2 | Pax8 | PEPCK Cre | mT/mG |
|---|---|---|---|---|
| WILD TYPE (WT) | *wt/wt* | *wt/wt* | +/Y | +/+ |
| MUTANT (MUT) | *fl/fl* | *fl/fl* | +/Y | +/+ |

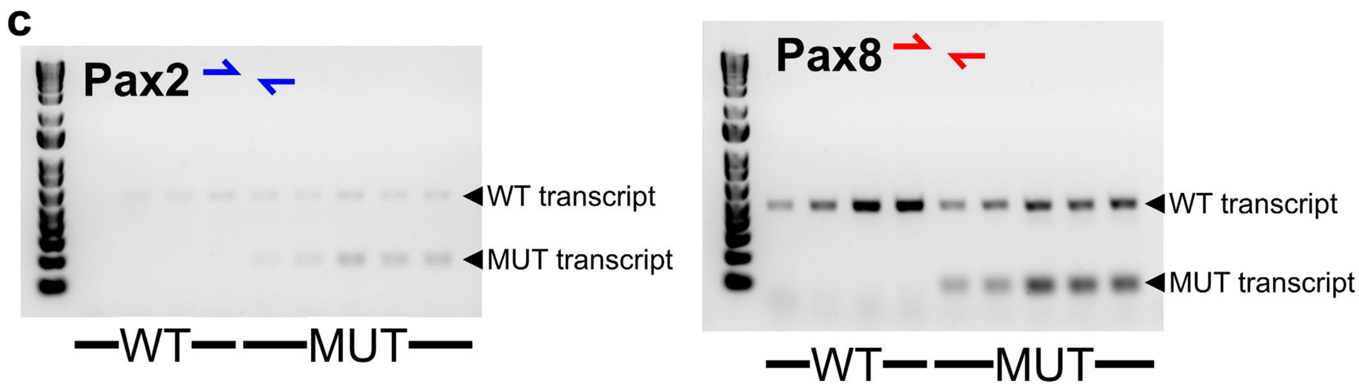


**Supplementary Figure S1. Detailed overview of mouse strain genetics.** (**a**) Locations of LoxP sites in the mutant (floxed) Pax2 and Pax8 genes. LoxP mediated recombination in mutant animals (MUT) resulted in loss of key exons in the DNA binding domain. Blue and red arrows show primer locations used for RT-PCR for Pax2 and Pax8, respectively (Supplementary Table S3). (**b**) Overview of strain abbreviations and animal genetics (*wt*: wild type Pax2 or Pax8 allele, *fl*: LoxP modified allele, +: presence of the transgene allele). (**c**) Semiquantitative RT-PCR (30 cycles) of cDNA obtained from lotus lectin purified proximal tubule cells of uninjured kidneys using primers shown in Supplementary Table S3. Similar amounts of both mutant and wild type transcript indicate heterogeneous recombination consistent with the PEPCK Cre activity shown in Figure 1a.

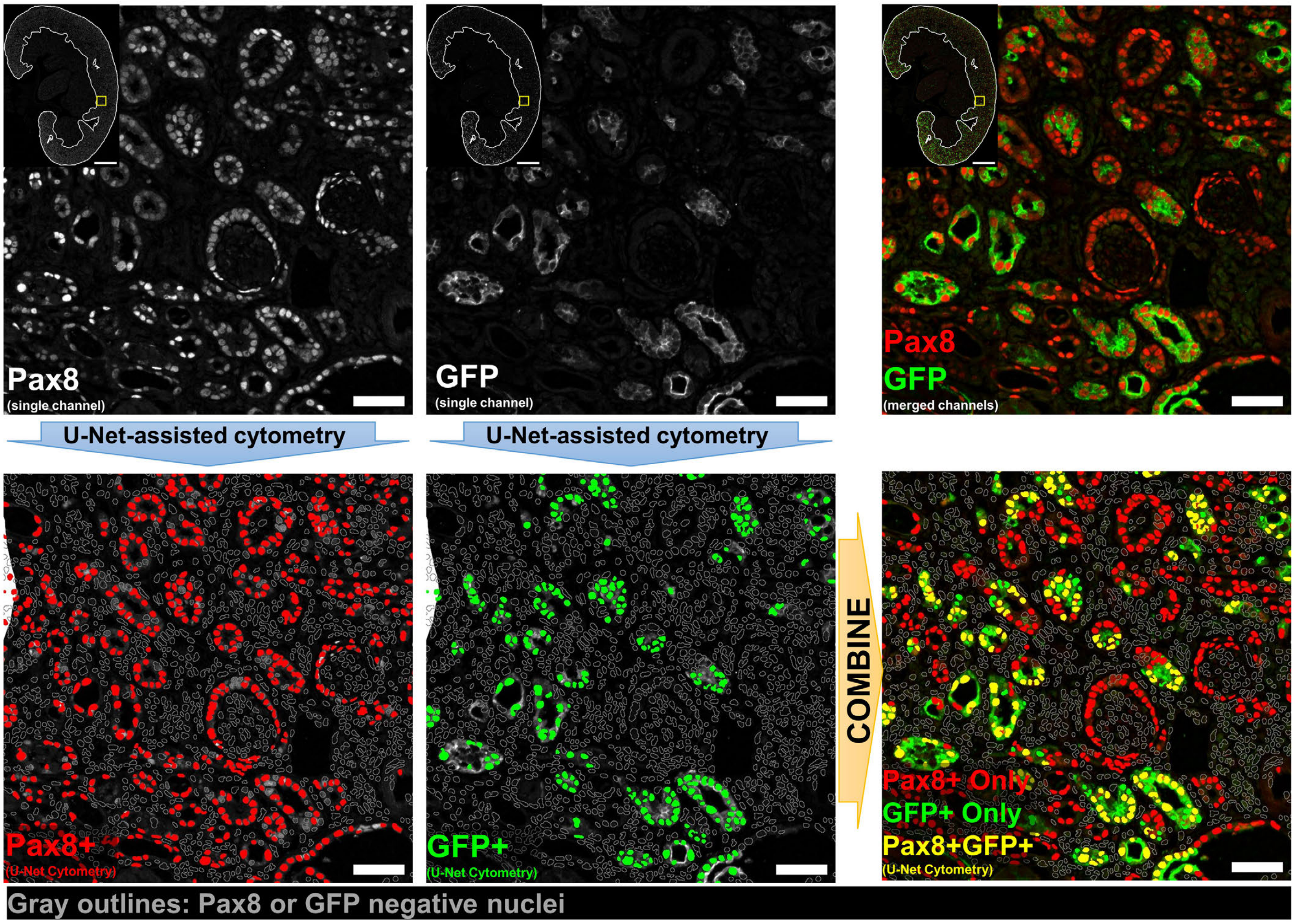


**Supplementary Figure S2. Overview of deep-learning-assisted tissue cytometry to measure proximal tubule cell Pax protein expression after injury**. Example images using Pax8 and GFP from an ischemic kidney corresponding to data shown in Figure 1d are shown. For clarity only one 400 µm square region is shown (yellow outline on full size image) but analysis was performed over the entire cortex and OSOM (white outline on full size images) for one section from each animal, typically ~2 x $10^5$ nuclei per section. Single channel images for markers of interest were segmented using a U-Net convolutional neural network specifically trained to identify positive cells, including their nuclei[27,52]. Nuclei were identified by segmenting a DAPI channel in each image (not shown). Each nucleus was classified based on the degree of overlap with the single channel segmentations (bottom row). Dual positive cells were determined by the intersection of the sets of single-marker-positive nuclei (bottom right). Gray outlines show additional nuclei analyzed but classified as negative for the marker(s) indicated. Scale bars: 1 mm (full section), 50 µm (region of interest).

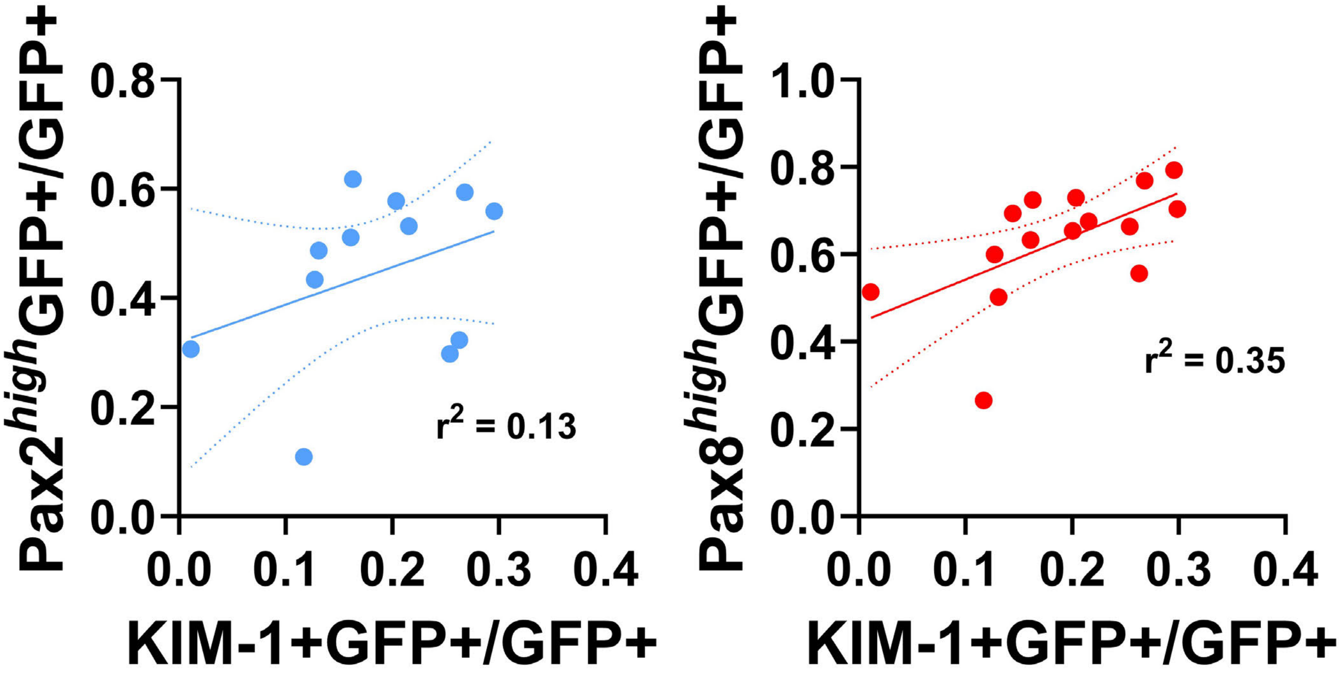


**Supplementary Figure S3. Correlation of Pax2 or Pax8 with KIM-1 expression in GFP+ proximal tubule cells**. Each point represents measurements from the same animal but from separate sections, as in Figure 1e (Pearson R = 0.36, p = 0.25 for Pax2, and Pearson R = 0.59, p = 0.02 for Pax8).

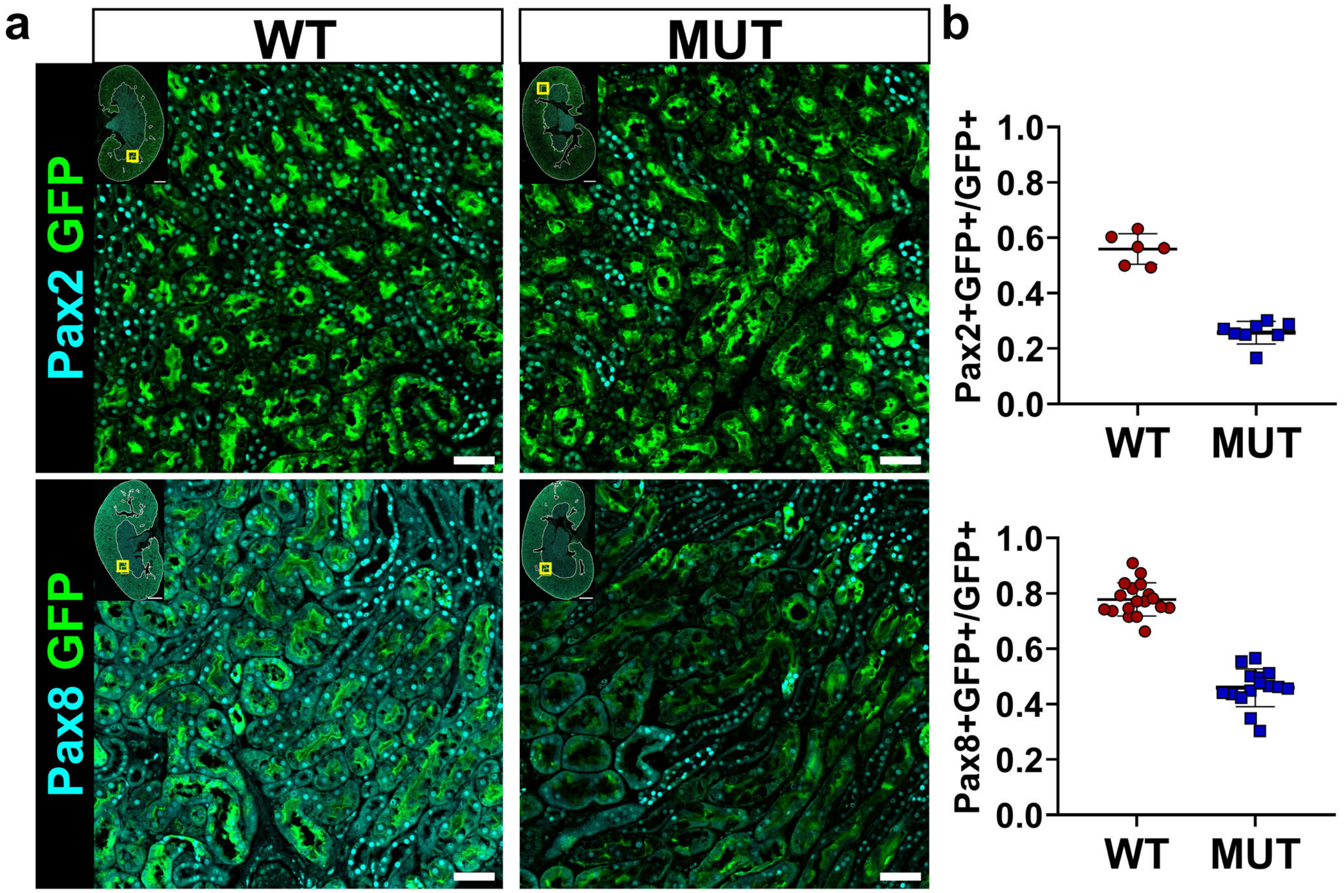


**Supplementary Figure S4. Pax2 and Pax8 are depleted in GFP expressing cells of mutant mice.** (**a**) Representative images showing Pax2 (top, cyan) and Pax8 (bottom, cyan) co-stained for GFP (green). Yellow box in inset shows image location in the larger section, white outline shows the cortex + OSOM used for analysis. Corresponding images showing Pax2 and Pax8 with villin are shown in Figure 2a. Scale bars: 1 mm (full section), 50 μm (region of interest). (**b**) Deep-learning-assisted quantification of Pax2 and Pax8 protein expression in GFP+ cells in the cortex + OSOM (*: $P < 0.0001$, t-test, $N = 6$-18 animals). Strain abbreviations-WT: $Pax2^{wt/wt}$, $Pax8^{wt/wt}$, PEPCK $Cre^{+/y}$, $mT/mG^{+/+}$; MUT: $Pax2^{fl/fl}$, $Pax8^{fl/fl}$, PEPCK $Cre^{+/y}$, $mT/mG^{+/+}$.

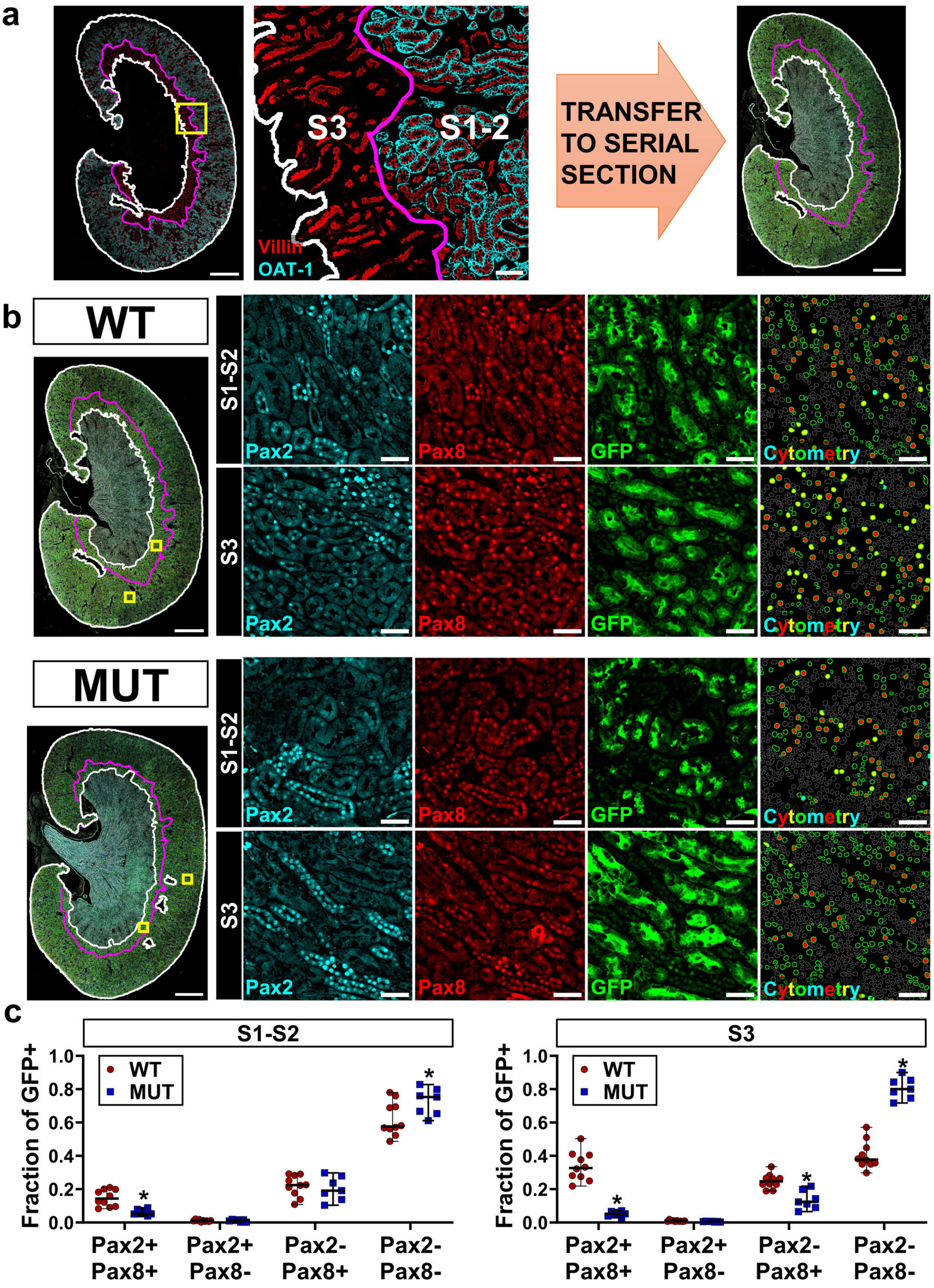


**Supplementary Figure S5. Both Pax2 and Pax8 are depleted in S3 proximal tubule.**
**(continued next page)**

**Supplementary Figure S5. (continued)** (**a**) Within the cortex + OSOM (white) The histologic border between S1-S2 and S3 (magenta) was defined by staining for villin (red, all proximal tubules) and organic anion transporter-1 (OAT-1, cyan, S2). These borders were manually transferred to a serial section to define compartments that were predominantly S1-S2 or S3. (**b**) Dual labeling of Pax2 and Pax8 (with GFP) was performed as outlined in the supplementary methods. Note that staining sensitivity was reduced with these methods, resulting in a lower fraction of detected nuclei compared with Figure 2 and Supplementary Figure S4. The fraction of GFP+ cells co-expressing Pax2 and Pax8 was quantified using deep-learning-assisted tissue cytometry throughout the entirety of each compartment. Representative regions are shown. Green outlines in the cytometry panel represent GFP+ cells analyzed. Fill represents detection of Pax2 (cyan), Pax8 (red), or both (yellow). Gray outlines are GFP- cells not included in analysis. Scale bars: 1 mm (full section), 50 μm (region of interest). (**c**) Quantification of GFP+ cells co-expressing Pax2 and Pax8 in each compartment (*: $P < 0.03$, two-way ANOVA with Sidak's post-hoc test, $N = 7\text{-}10$ animals). Strain abbreviations-WT: $Pax2^{wt/wt}$, $Pax8^{wt/wt}$, PEPCK $Cre^{+/y}$, $mT/mG^{+/+}$; MUT: $Pax2^{fl/fl}$, $Pax8^{fl/fl}$, PEPCK $Cre^{+/y}$, $mT/mG^{+/+}$.

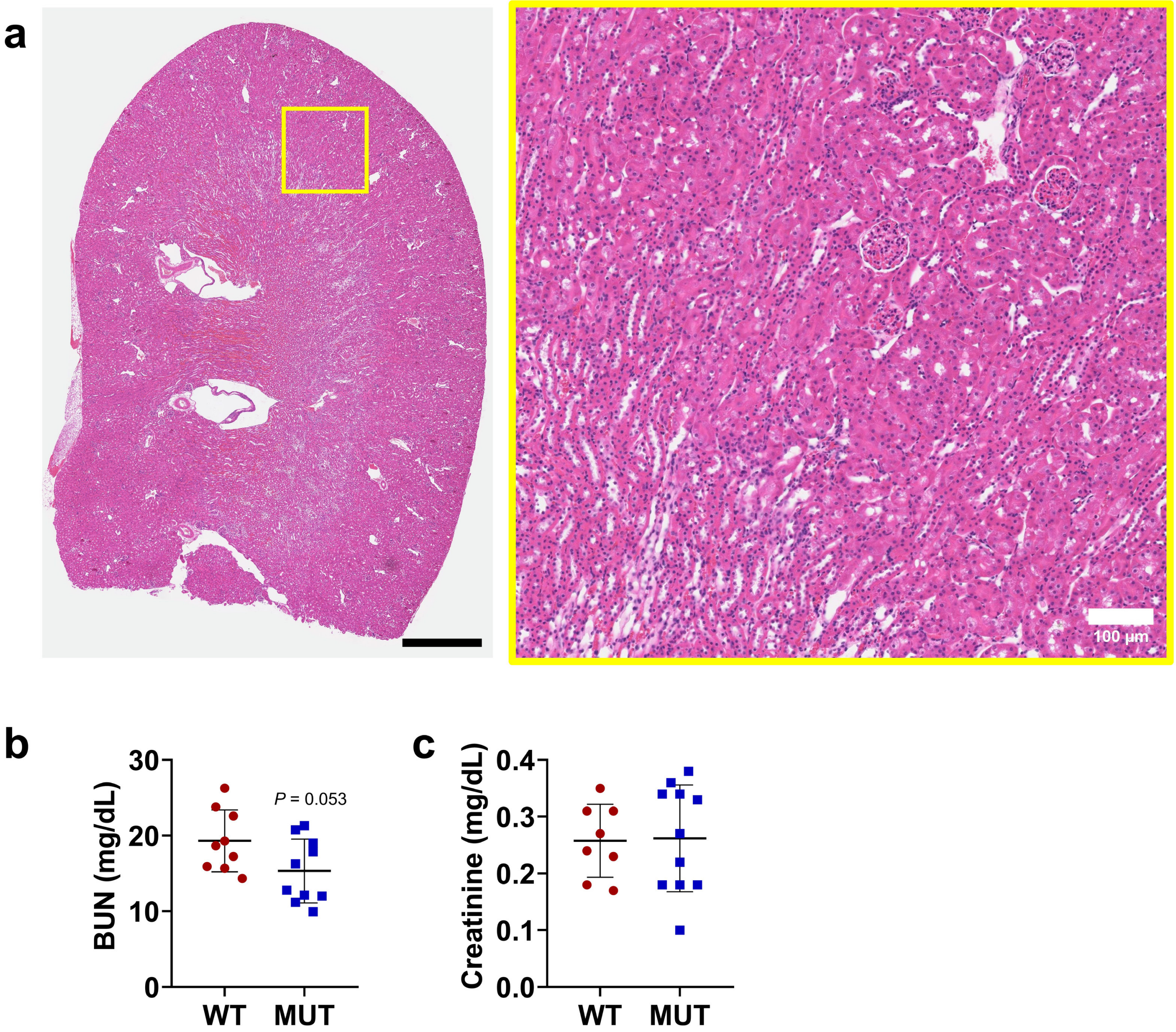


**Supplementary Figure S6. Mutant animals show normal gross histology and similar kidney function compared with WT controls**. (**a**) Representative area of hematoxylin and eosin-stained kidney at the corticomedullary junction. Note, the lower pole of this sample was intentionally excised for PCR genotype confirmation. Scale bars: 1 mm whole kidney, 100 µm inset. (**b**) Serum blood urea nitrogen (BUN) and (**c**) creatinine measured in 12-week-old uninjured mice. Strain abbreviations-WT: $Pax2^{wt/wt}$, $Pax8^{wt/wt}$, PEPCK $Cre^{+/y}$, $mT/mG^{+/+}$; MUT: $Pax2^{fl/fl}$, $Pax8^{fl/fl}$, PEPCK $Cre^{+/y}$, $mT/mG^{+/+}$.

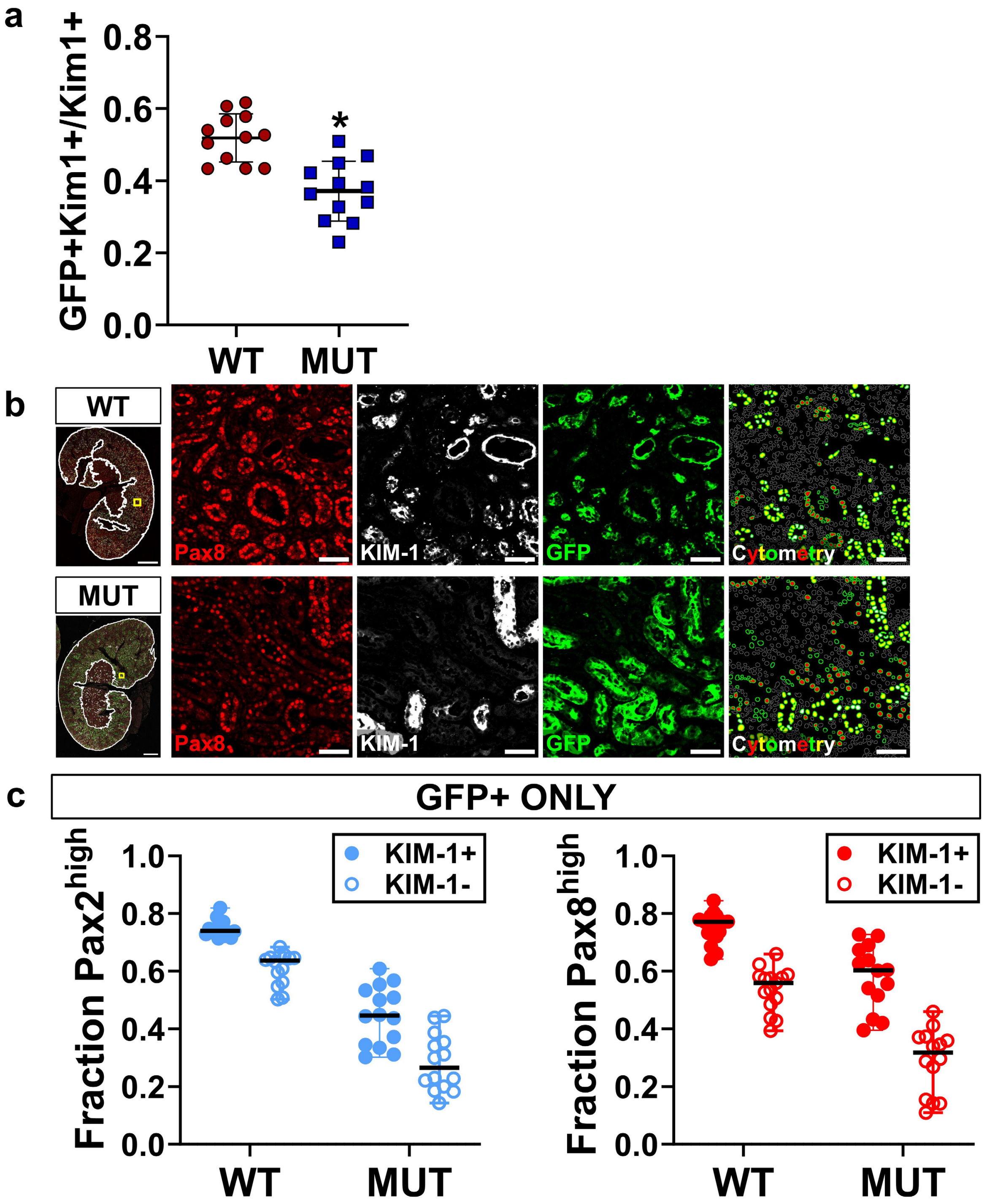


**Supplementary Figure S7. Pax2 and Pax8 mutant cells are less likely to contribute to incomplete repair marked by KIM-1**. (**a**) Quantification of the fraction of GFP+ cells in the subpopulation of KIM-1+ cells, which marks incomplete repair, analyzed as for Vcam1 in Figure 4. Measurements were made across the cortex + OSOM for each section (*: $P < 0.0001$, t-test, $N = 12$ animals). (**b**) Pax2 or Pax8, KIM-1, and GFP expression was quantified using deep-learning-assisted tissue cytometry in the cortex + OSOM. Green outlines in the cytometry panel represent GFP+ cells analyzed. Fill represents detection of Pax8 (red), KIM-1 (white), or both (yellow). Representative regions are shown for Pax8. An identical analysis was performed for Pax2. (**c**) The fraction of Pax2 or Pax8 in the GFP+ population grouped by KIM-1 status (all comparisons were significant, $P \leq 0.0001$, except WT/KIM-1- vs MUT/KIM-1+ for Pax8, two-way ANOVA with Sidak's post-hoc test, $N = 14\text{-}15$ animals). Strain abbreviations-WT: $Pax2^{wt/wt}$, $Pax8^{wt/wt}$, PEPCK $Cre^{+/y}$, $mT/mG^{+/+}$; MUT: $Pax2^{fl/fl}$, $Pax8^{fl/fl}$, PEPCK $Cre^{+/y}$, $mT/mG^{+/+}$.

**a**

| Strain Abbreviation | Pax2 | Pax8 | PEPCK Cre | mT/mG |
|---|---|---|---|---|
| ALTERNATIVE WILD TYPE ($WT_o$) | *fl/fl* | *fl/fl* | *-/Y* | *-/-* |
| ALTERNATIVE MUTANT ($MUT_o$) | *fl/fl* | *fl/fl* | *+/Y* | *-/-* |

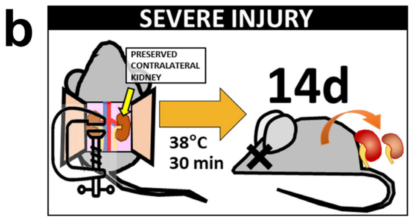


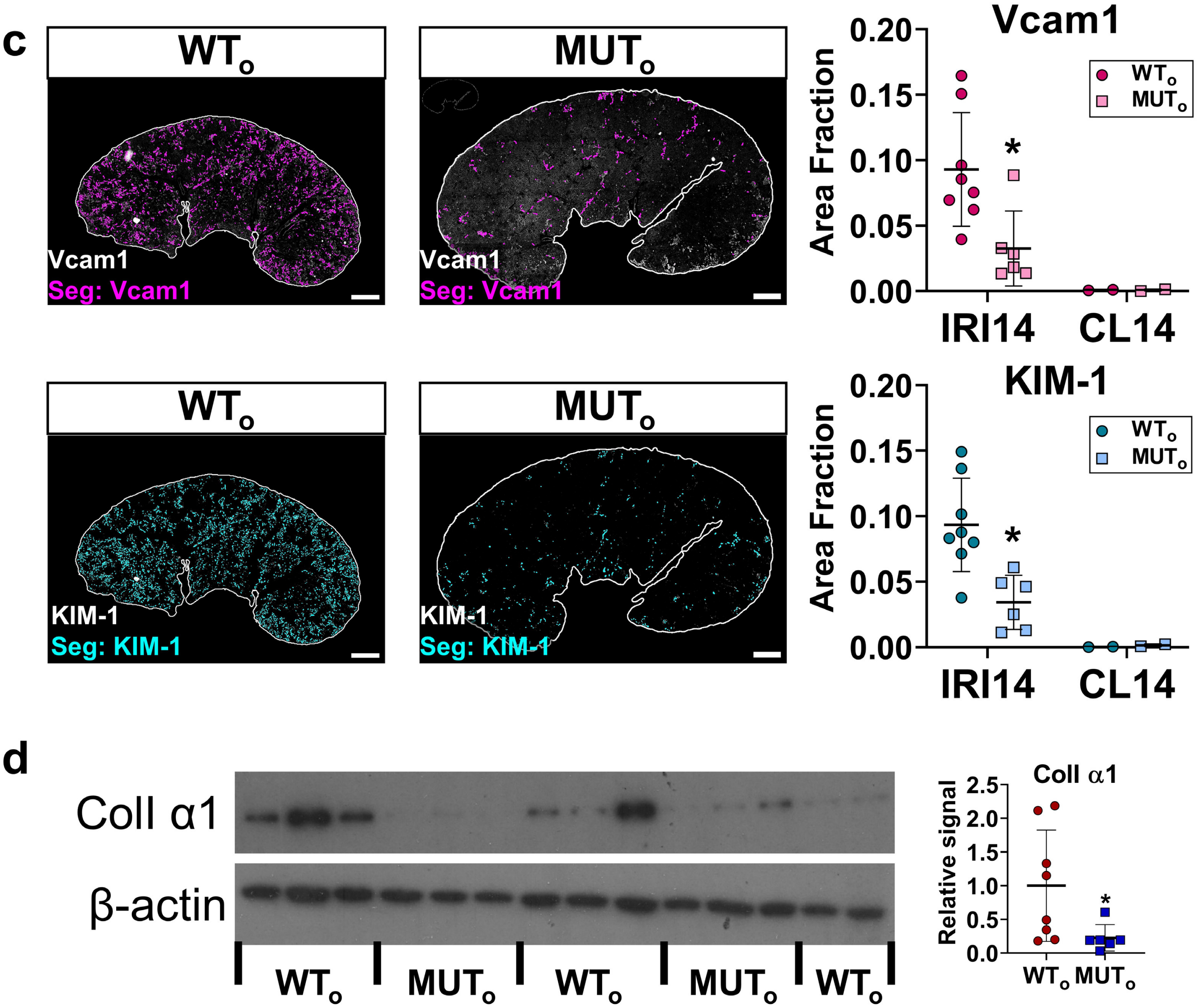


**Supplementary Figure S8. Pax2 and Pax8 mutant mice with alternative genetics are also protected from chronic injury after severe IRI.** (**a**) Genetics of alternative mice strains where $WT_o$ and $MUT_o$ status was determined by the absence or presence of PEPCK Cre (all with animals with $Pax2^{fl/fl}$, $Pax8^{fl/fl}$) rather than the absence or presence of floxed Pax2 and Pax8 alleles. The mT/mG allele (GFP reporter) was excluded because its expression would also differ between these strains. (**b**) Animals were subjected to severe IRI with intact contralateral kidney as in Figure 3. Scale bars: 1 mm. (**c**) Representative images, segmentation outputs, and quantification of segmented area fraction for Vcam1 and KIM-1 (*: $P$ = 0.008 and 0.002, respectively, t-test for IRI14 only, $N$ = 6-8 animals). (**d**) Western blot for Collagen I α1 and densitometry quantification (*: $P$ = 0.03, t-test, $N$ = 6-8 animals).

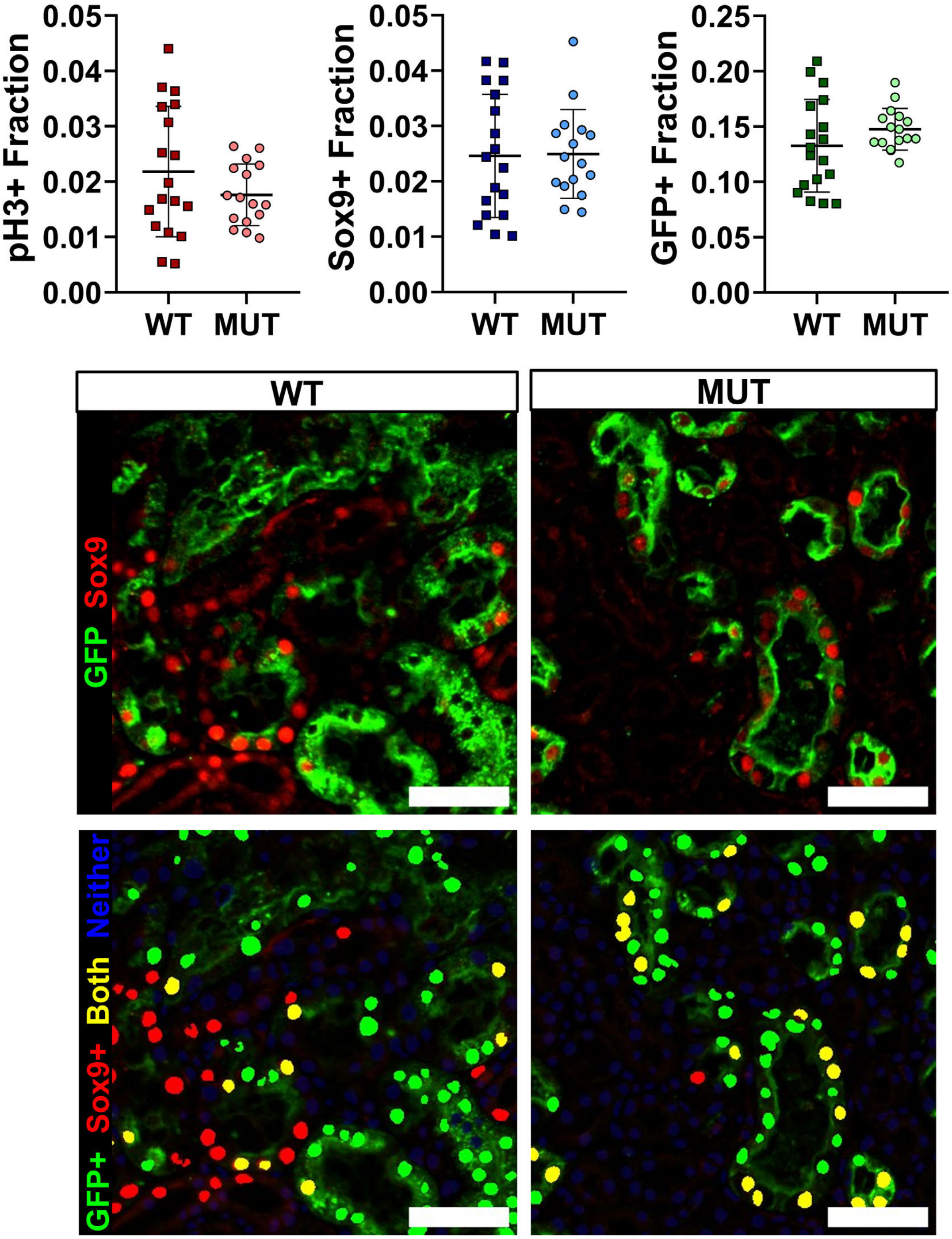


**Supplementary Figure S9. Supporting data for Figure 5.** (**a**) Fraction of all cells detected by U-Net assisted cytometry for pH3, Sox9, and GFP in samples 2 d after IRI. No differences were detected (p = 0.19, 0.92 and 0.18 respectively, t-test, *N* = 16-18 animals) (**b**) Original images (top) and representative images showing tissue cytometry (bottom) for GFP and Sox9 showing GFP+ (green), Sox9+ (red), dual positive (yellow) or negative (blue) cells 2 d after IRI. The background was dimmed in the cytometry images for clarity. Scale bars: 50 μm. Strain abbreviations-WT: Pax2$^{wt/wt}$, Pax8$^{wt/wt}$, PEPCK Cre$^{+/y}$, mT/mG$^{+/+}$; MUT: Pax2$^{fl/fl}$, Pax8$^{fl/fl}$, PEPCK Cre$^{+/y}$, mT/mG$^{+/+}$.

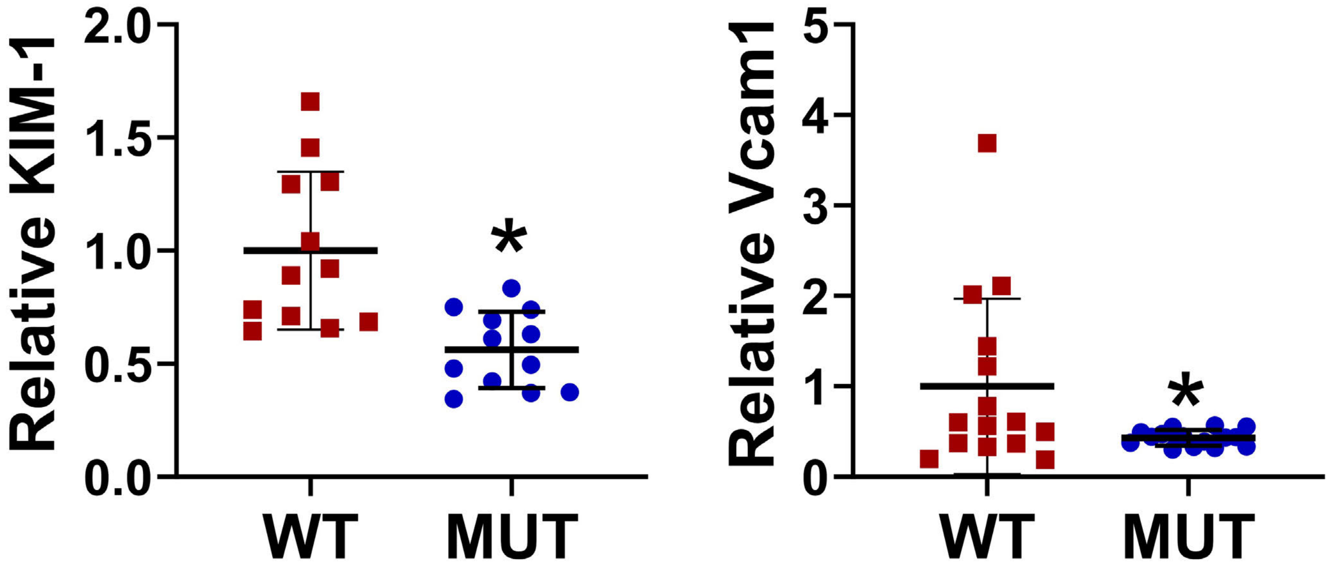


**Supplementary Figure S10. Pax2 and Pax8 mutant mice develop less chronic injury after IRI.** RNA expression of KIM-1 and Vcam1 was measured in whole kidney isolates 28 d after IRI (*: $P$ = 0.0008 and 0.04, respectively, t-test, $N$ = 15 animals)**.** Strain abbreviations-WT: $Pax2^{wt/wt}$, $Pax8^{wt/wt}$, PEPCK $Cre^{+/y}$, $mT/mG^{+/+}$; MUT: $Pax2^{fl/fl}$, $Pax8^{fl/fl}$, PEPCK $Cre^{+/y}$, $mT/mG^{+/+}$.

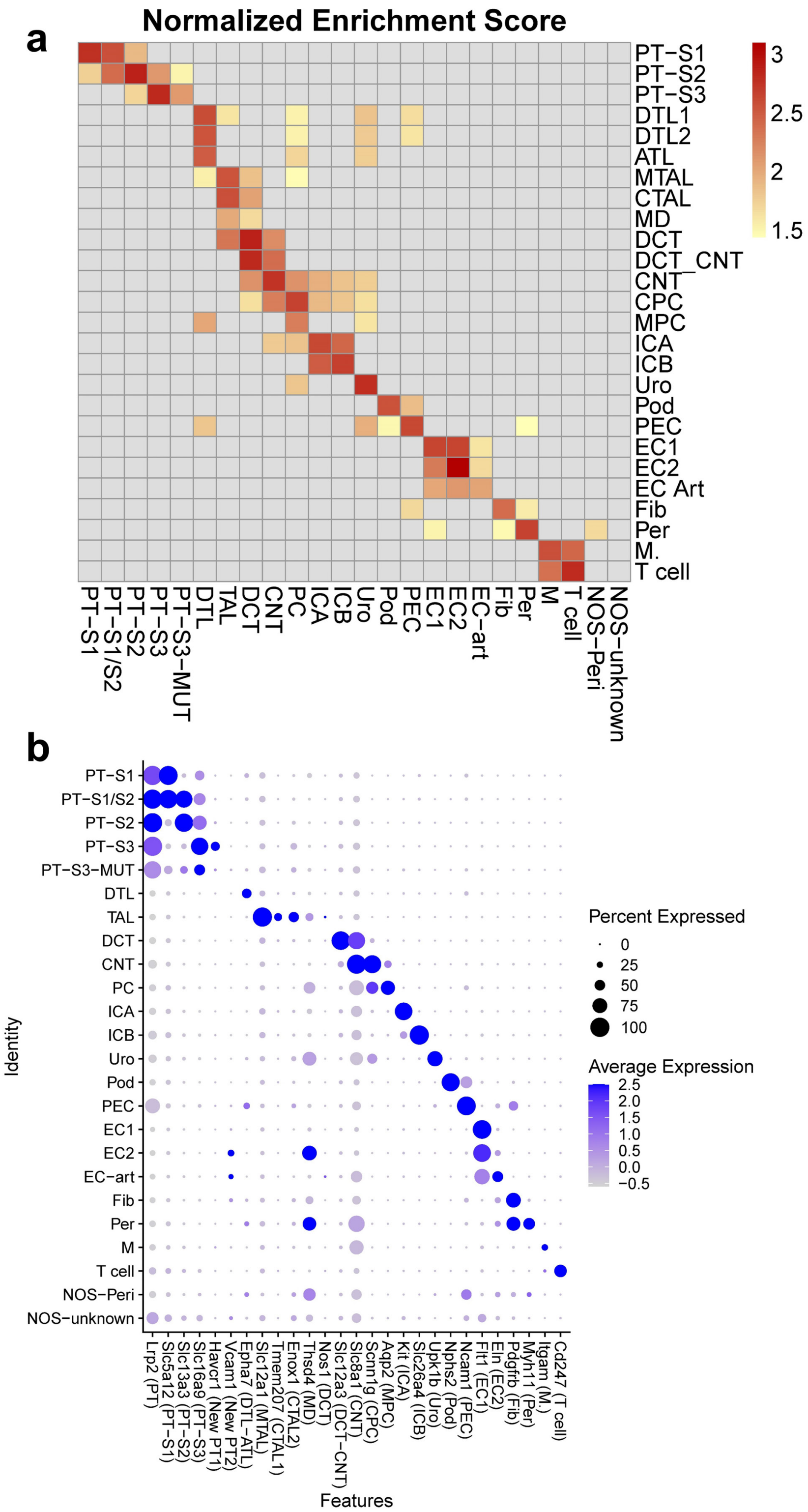


**Supplementary Figure S11. Cluster alignment of whole kidney snRNA seq.** (**a**) Heat map of normalized enrichment scores obtained from GSEA comparing differentially expressed genes in each new cluster (upregulated relative to all other clusters) with the top 50 upregulated genes in each cluster from Kirita et al[6]. Only enrichments with adjusted p-value < 0.05 are colored. Comparisons without significant enrichment are shown in gray. GSEA results were used to assign cluster names in Figure 6a. (**b**) Dot plot of expression of example marker genes for each cluster, as identified in Kirita et al, in the clusters assigned in (**a**).

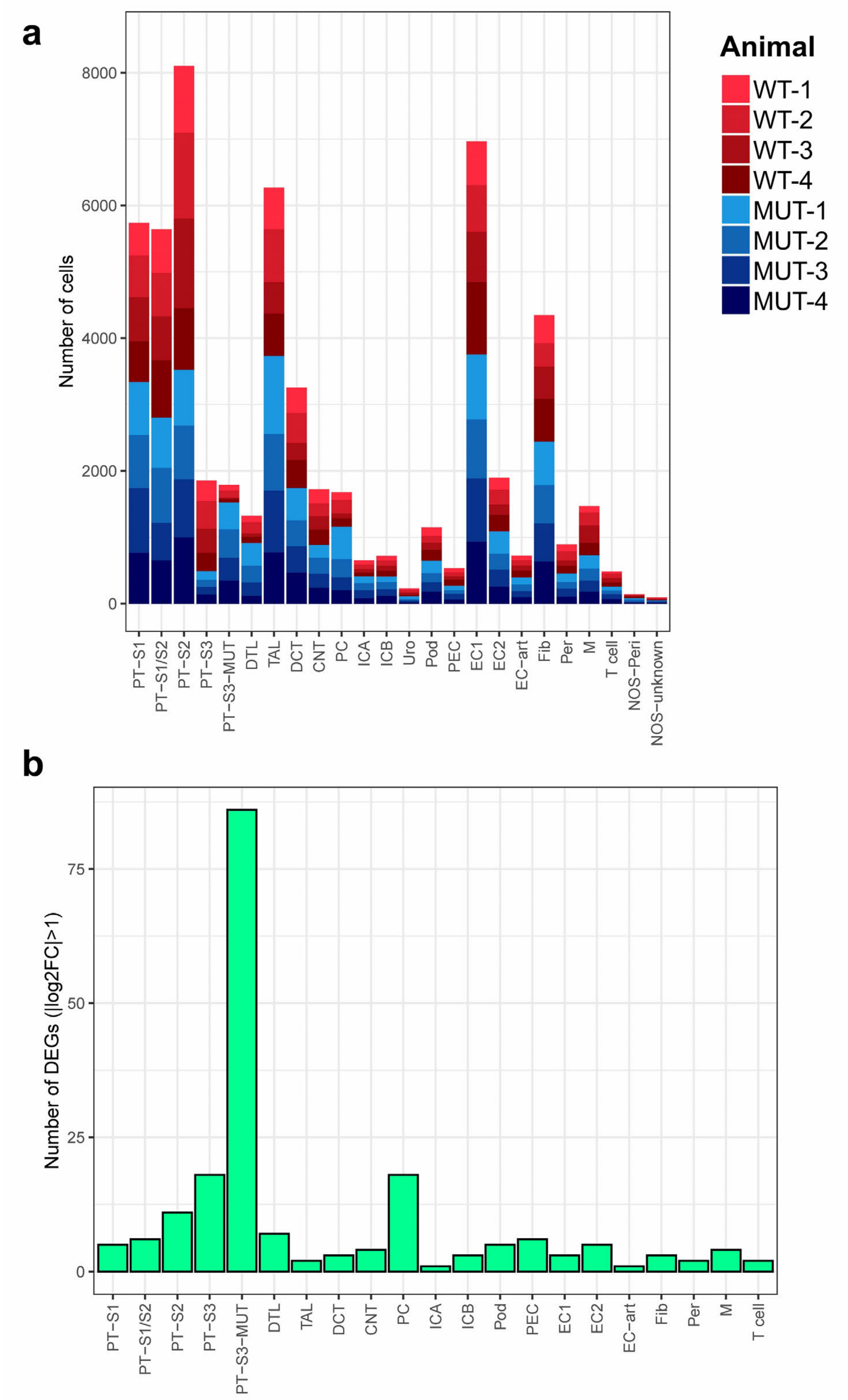


**Supplementary Figure S12. Cluster composition and differentially expressed genes from whole kidney snRNA seq.** (**a**) Histogram of cells assigned to each cluster. WT animals are shown in shades of red, MUT animals in shades of blue. (**b**) Number of differentially expressed genes (DEGs) between MUT and WT animals in each cluster with |log2 fold change| > 1. Strain abbreviations-WT: Pax2$^{wt/wt}$, Pax8$^{wt/wt}$, PEPCK Cre$^{+/y}$, mT/mG$^{+/+}$; MUT: Pax2$^{fl/fl}$, Pax8$^{fl/fl}$, PEPCK Cre$^{+/y}$, mT/mG$^{+/+}$.

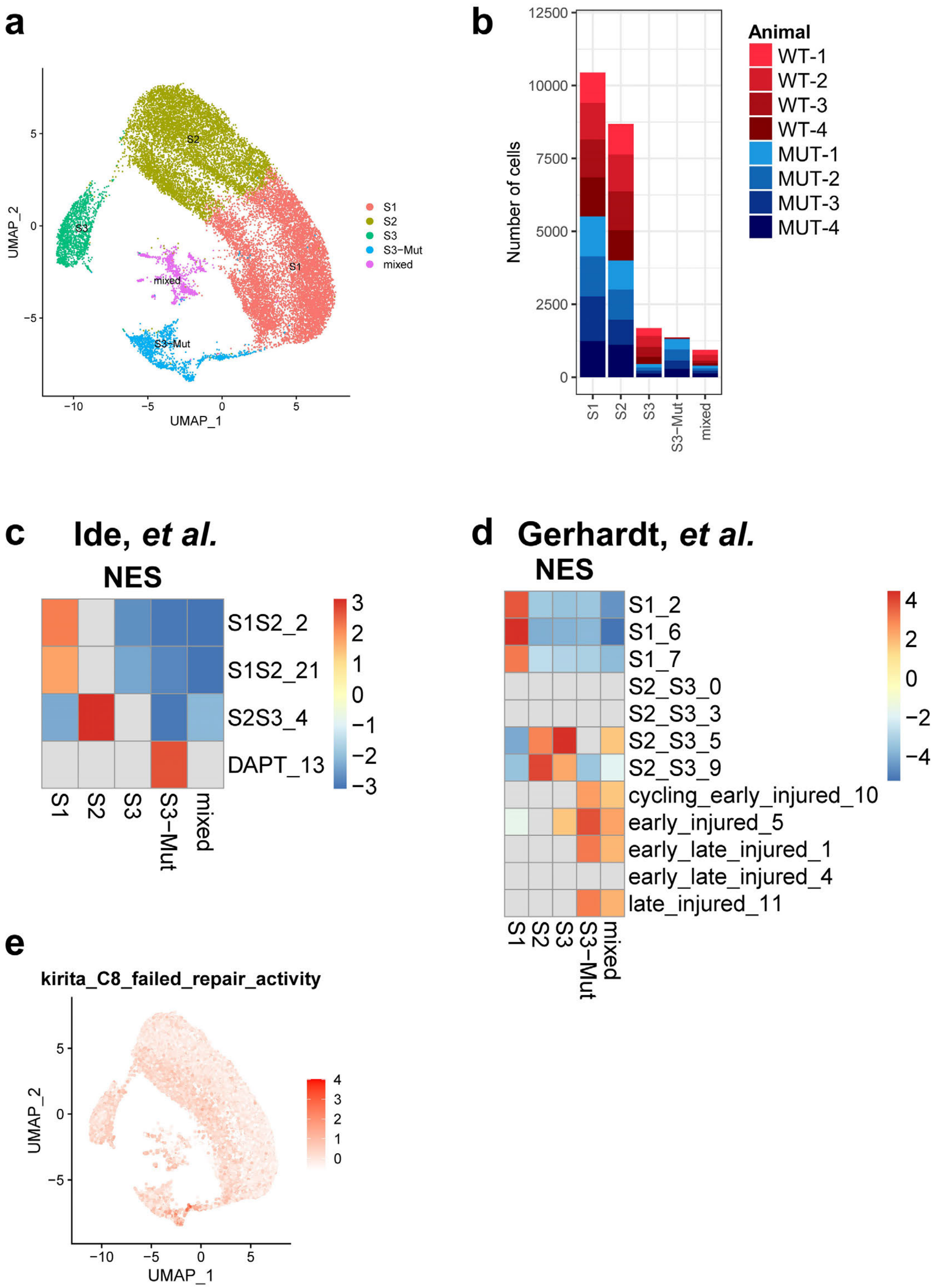


**Supplementary Figure S13. Composition of subclustered proximal tubule cells and gene set enrichment (GSEA) with alternative single cell/single nucleus RNA sequencing data sets.** (**a**) UMAP plot of subclustered proximal tubules (all cells). (**b**) Histogram of cells assigned to each cluster. WT animals are shown in shades of red, MUT animals in shades of blue. (**c**) GSEA of differentially expressed genes in the proximal tubule clusters shown in (**a**) compared with the top 50 differentially expressed genes in proximal tubule clusters in (**c**) Ide et al[13] and (**d**) Gerhardt et al[28]. (**e**) Average z-score projection of genes upregulated in "failed repair" proximal tubules[6]. Strain abbreviations-WT: $Pax2^{wt/wt}$, $Pax8^{wt/wt}$, PEPCK $Cre^{+/y}$, $mT/mG^{+/+}$; MUT: $Pax2^{fl/fl}$, $Pax8^{fl/fl}$, PEPCK $Cre^{+/y}$, $mT/mG^{+/+}$.

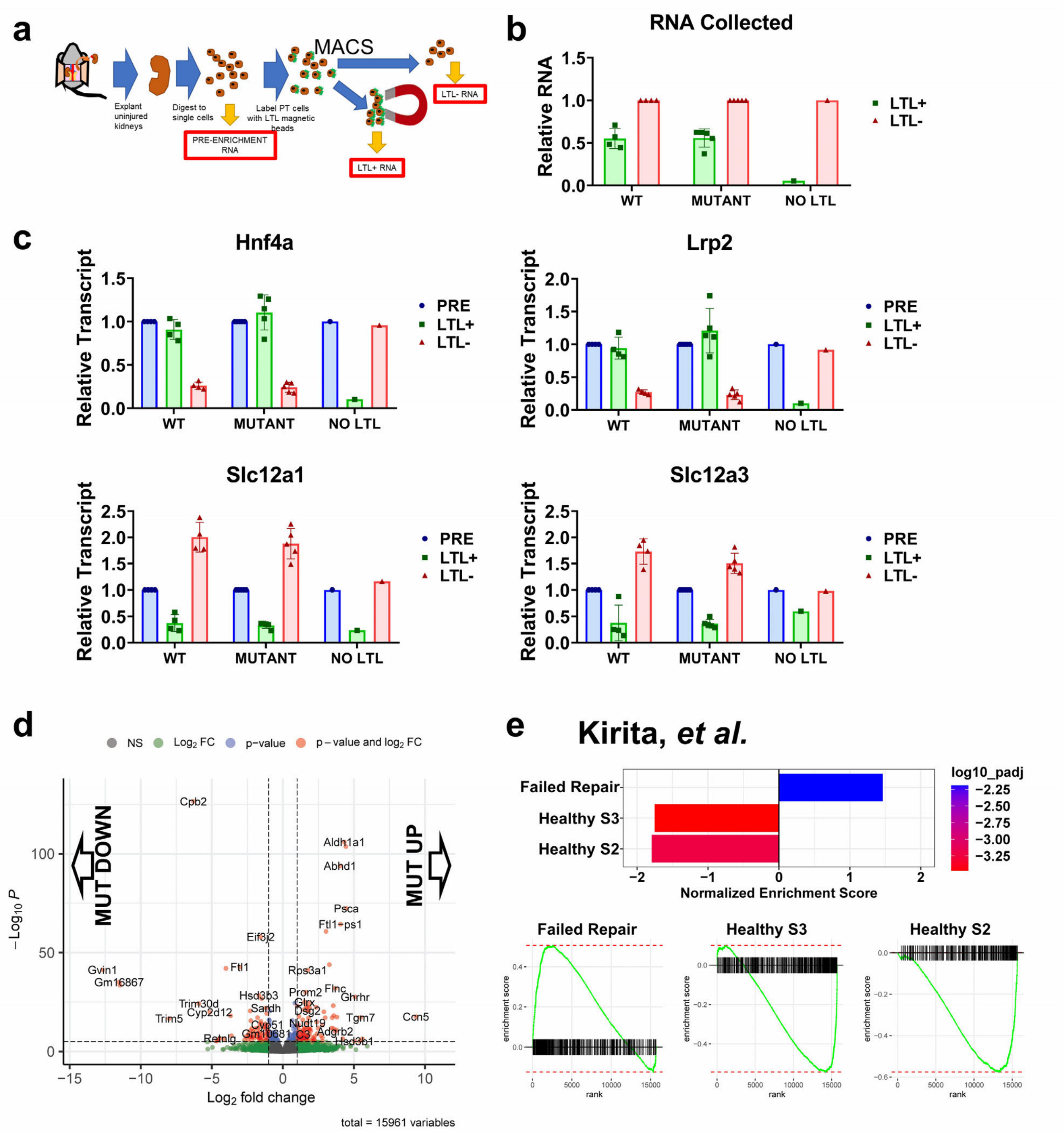


**Supplementary Figure S14. Purification and transcriptional profiling of proximal tubules using magnetic activated cell sorting (MACS)**. (**a**) Overview of proximal tubule (PT) cell purification using lotus tetragonolobus lectin (LTL) magnetic beads. (**b**) Total RNA collected from each of the LTL+ and LTL- fractions for each sample (1-5 kidneys). No LTL indicates a sample prepared identically except without addition of LTL. (**c**) Relative expression of proximal tubule (Hnf4a, Lrp2), thick ascending limb of the loop of Henle (Slc12a1), and distal tubule (Slc12a2) in the various fractions for each sample using quantitative RT-PCR (before sequencing). (**d**) Volcano plot of differentially expressed genes determined by bulk RNA seq of purified proximal tubule cells from MUT relative to WT animals (**e**) Gene set enrichment analysis for differentially expressed genes in MUT relative to WT compared with the differentially expressed genes in all proximal tubule clusters from Kirita et al[6]. Only GSEA results with adjusted *P* < 0.05 are shown. GSEA plots are shown below for each cluster. Strain abbreviations-WT: Pax2$^{wt/wt}$, Pax8$^{wt/wt}$, PEPCK Cre$^{+/y}$, mT/mG$^{+/+}$; MUT: Pax2$^{fl/fl}$, Pax8$^{fl/fl}$, PEPCK Cre$^{+/y}$, mT/mG$^{+/+}$.

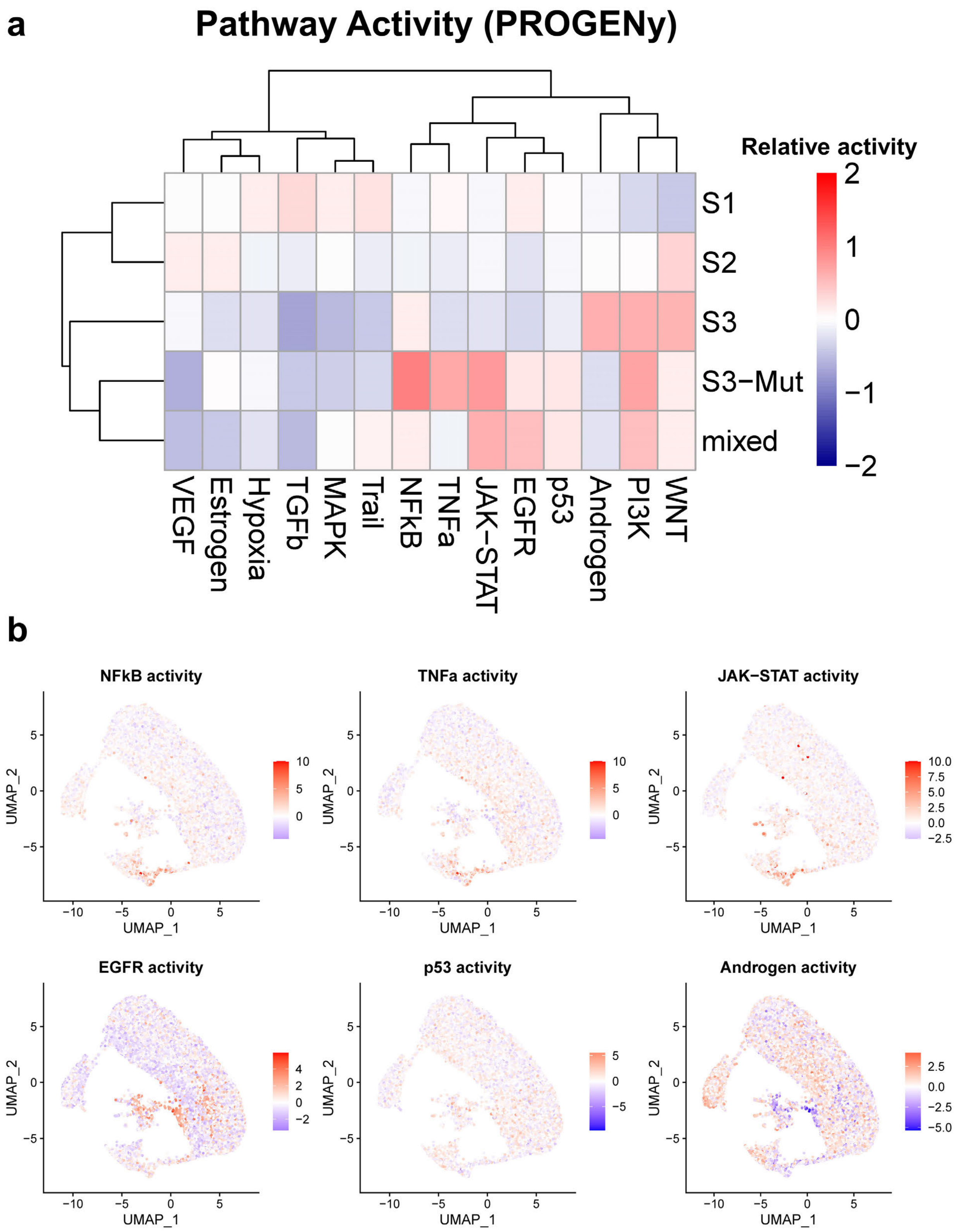


**Supplementary Figure S15. Proximal tubule pathway activation using PROGENy**. (**a**) Heatmap of relative pathway activation in proximal tubule clusters defined in Figure 6b. Activity was inferred using PROGENy[29-31] with decoupleR[53]. (**b**) Cell-wise pathway activity plotted in the UMAP space defined in Figure 6b for pathways with substantial differences between the healthy S3 (S3) and mutant S3 (S3-mut) clusters.

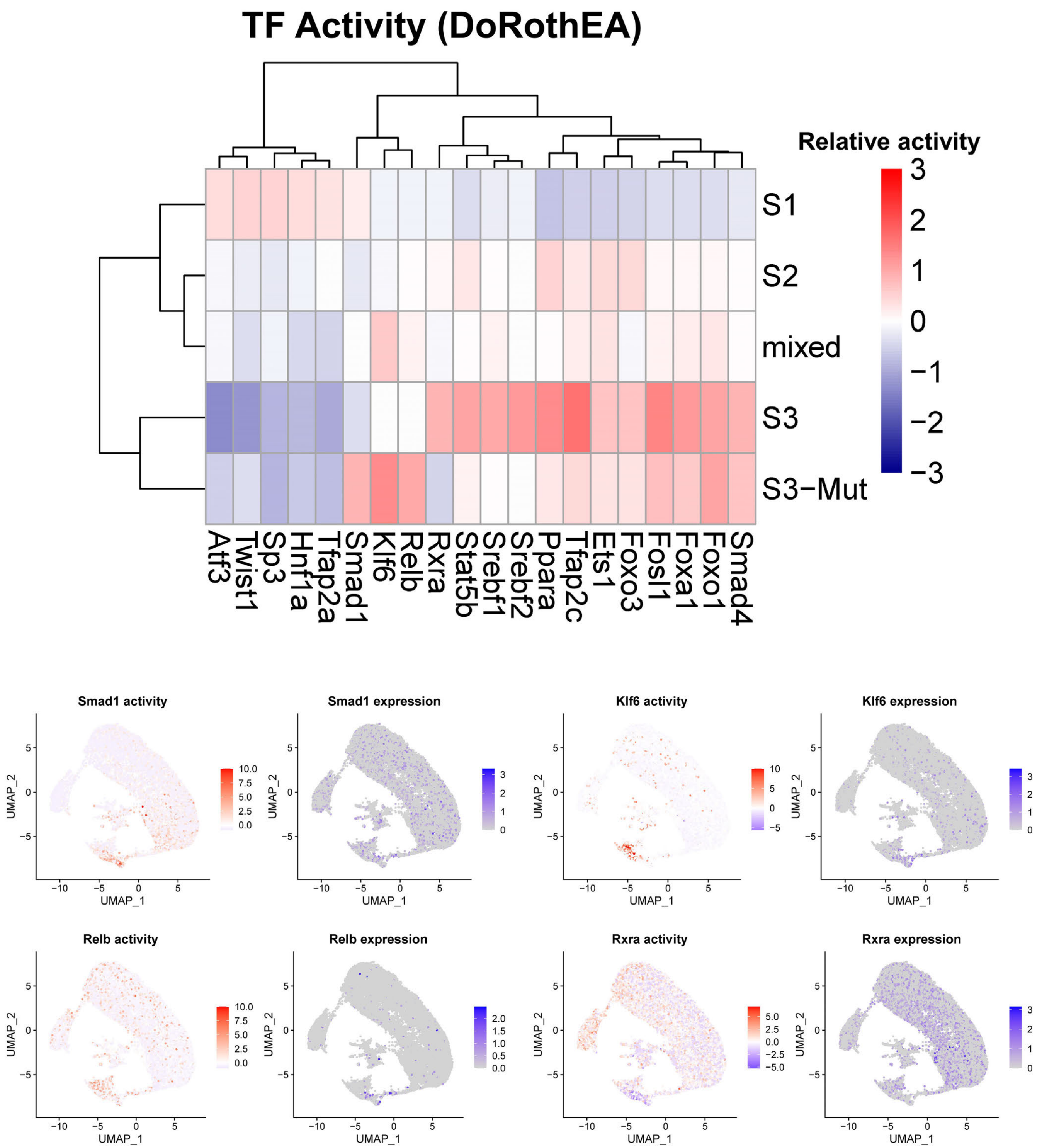


**Supplementary Figure S16. Transcription factor activity in proximal tubule clusters using DoRothEA**. (**a**) Heatmap of relative transcription factor activity for high confidence transcription factor signatures defined by DoRothEA[32] (level "A") for proximal tubule clusters defined in Figure 6b. The top 20 most variable signatures across clusters are shown. (**b**) Cell-wise transcription factor activity (red) and transcription factor expression (purple) plotted in the UMAP space defined in Figure 6b for transcription factor activities with substantial differences between the healthy S3 (S3) and mutant S3 (S3-mut) clusters.

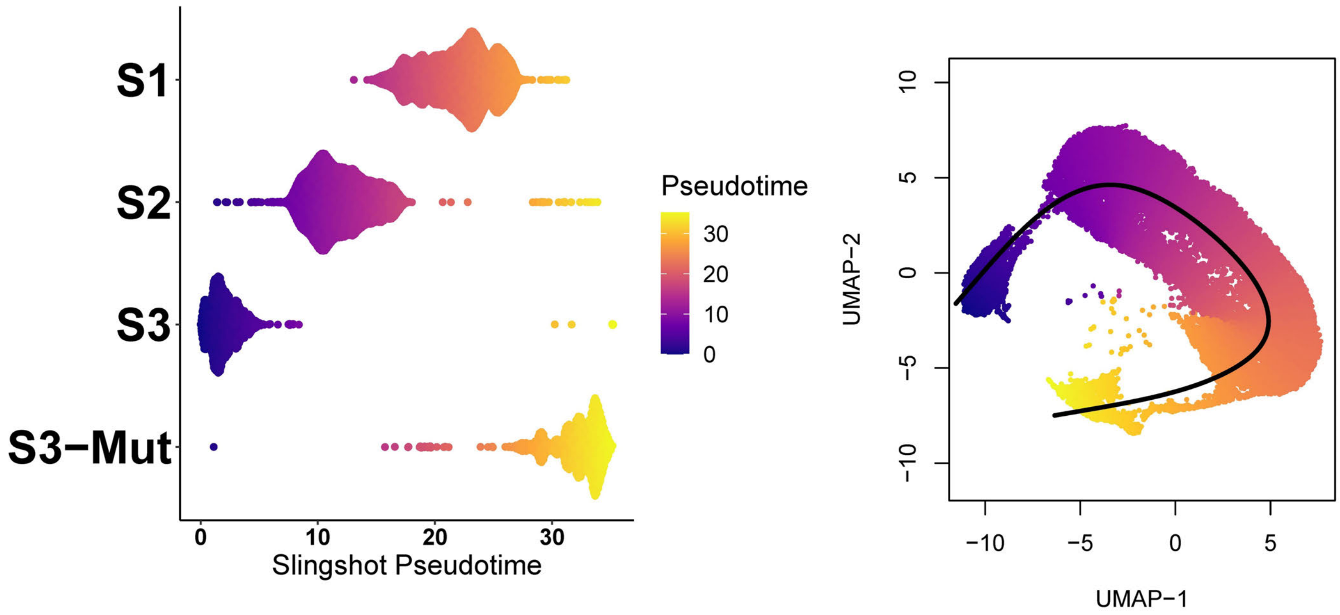


**Supplementary Figure S17. Pseudotime trajectory analysis of proximal tubule cells.** Pseudotime ordering was performed using Slingshot[35]. For clarity, the "Mixed" proximal tubule cluster was excluded from the analysis because it was small, derived equally from mutant and WT animals, and did not map to known proximal tubule cell types. See Supplementary Methods for details.

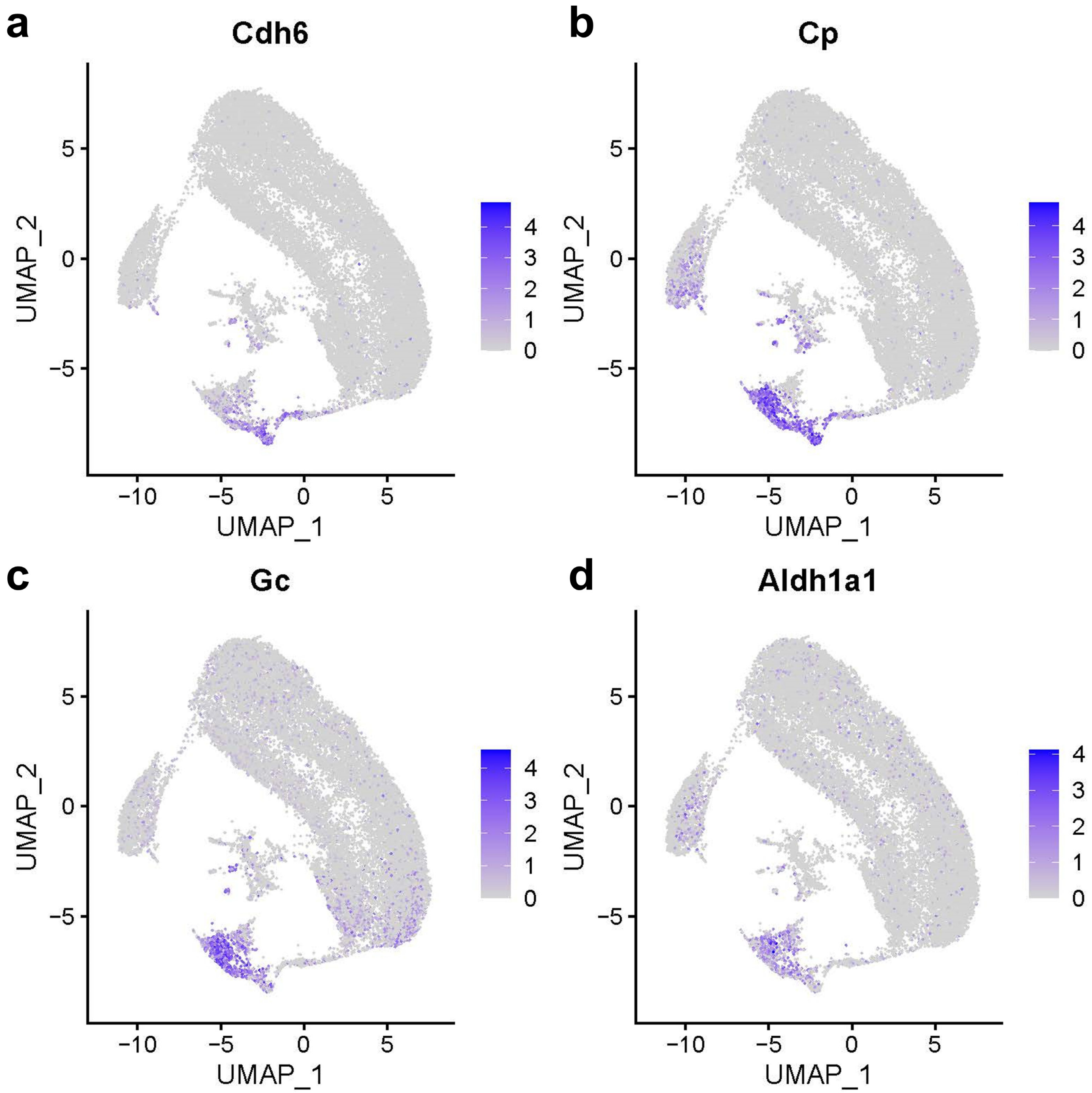


**Supplementary Figure S18. Expression of selected markers of mutant S3 proximal tubule by scRNA seq.** Feature plots for the cell-level expression patterns for Cdh6 (**a**), Cp (**b**), Gc (**c**), and Aldh1a1 (**d**).

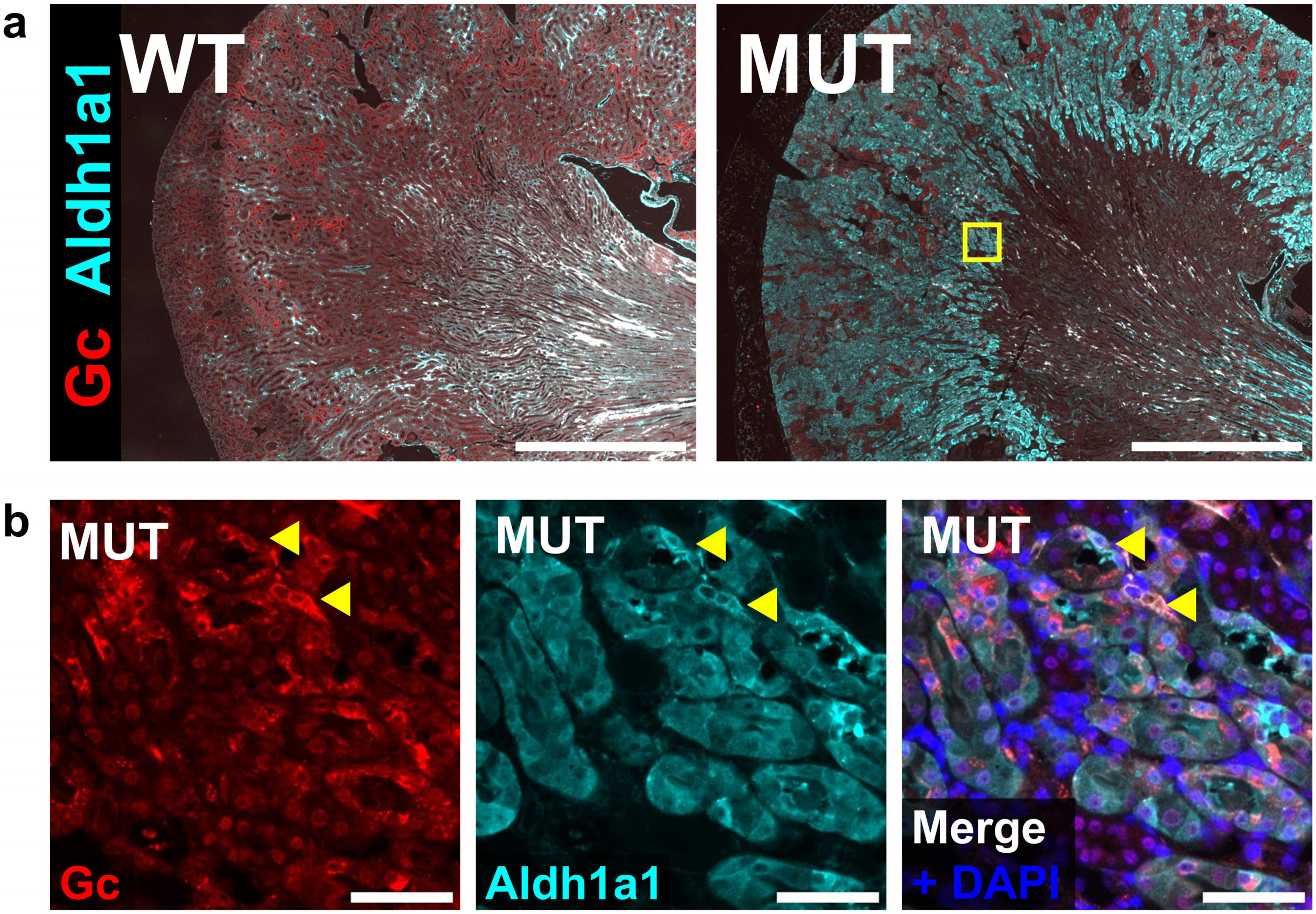


**Supplementary Figure S19. Expression of alternative markers of mutant S3 proximal tubule by immunofluorescence.** (**a**) Representative wide field images of Gc and Aldh1a1 staining of WT and MUT animals with equivalent acquisition settings. Scale bar: 1 mm. (**b**) Representative region in the S3 segment showing focal increased Gc and Aldh1a1 staining (yellow arrowheads). Scale bar: 50 µm. Strain abbreviations-WT: $Pax2^{wt/wt}$, $Pax8^{wt/wt}$, PEPCK $Cre^{+/y}$, $mT/mG^{+/+}$; MUT: $Pax2^{fl/fl}$, $Pax8^{fl/fl}$, PEPCK $Cre^{+/y}$, $mT/mG^{+/+}$.

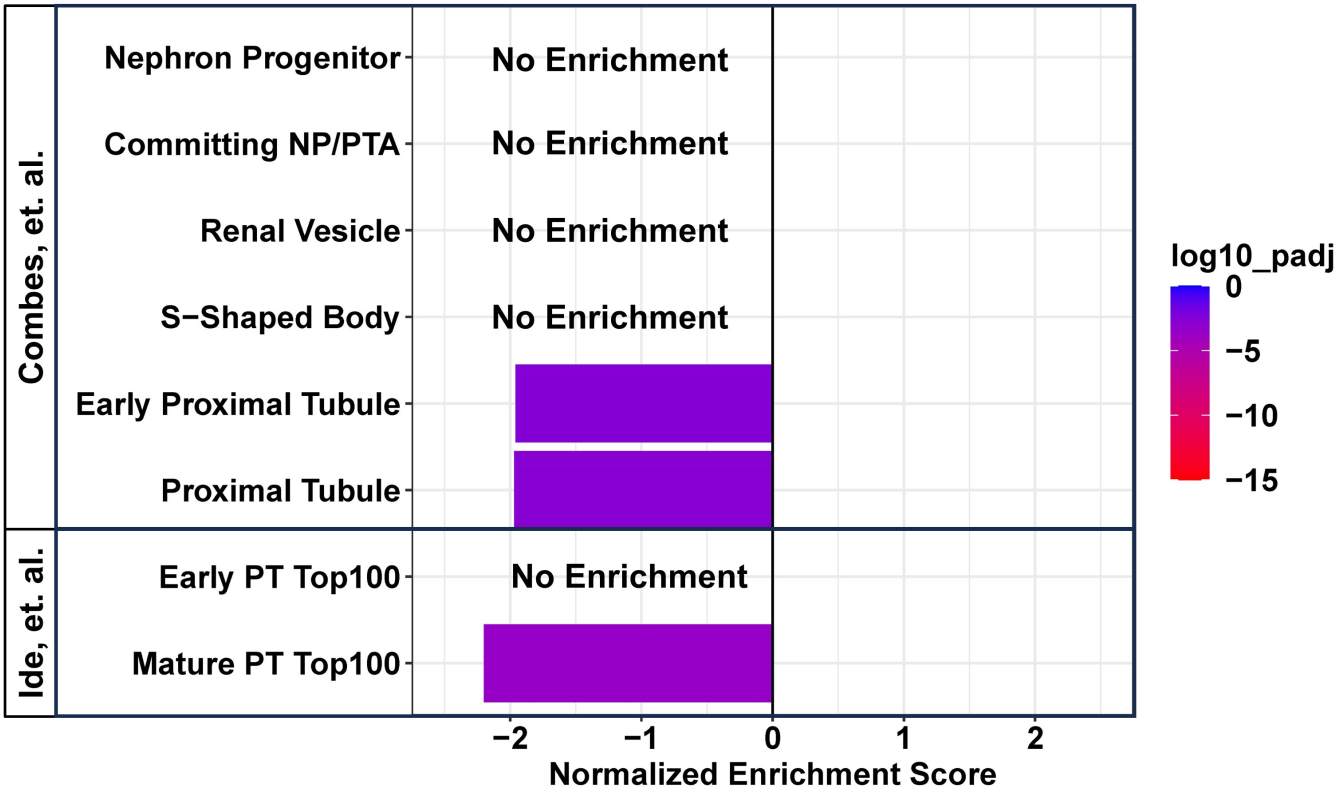


**Supplementary Figure S20. Mutant S3 cells do not show positive enrichment of developmental markers.** Gene set enrichment analysis for differentially expressed genes in S3-Mut relative to normal S3 compared with the differentially expressed genes in developing nephrons as described in Combes, et al[38] and Ide, et al[13,37]. Only GSEA results with adjusted *P* < 0.05 are shown.

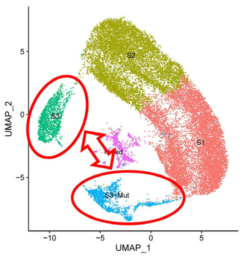


| UP REGULATED GENES | | | | |
|---|---|---|---|---|
| Gene | Female | HP | CR | Product name |
| Cpn1 | ■ | ■ | ■ | carboxypeptidase N, polypeptide 1 |
| Gsta2 | ■ | ■ | ■ | glutathione S-transferase, alpha 2 (Yc2) |
| Cdkl1 | ■ | ■ | | cyclin-dependent kinase-like 1 (CDC2-related kinase) |
| Gc | ■ | | ■ | vitamin D binding protein |
| Gsta3 | ■ | | ■ | glutathione S-transferase, alpha 3 |
| Cyp2d26 | ■ | | ■ | cytochrome P450, family 2, subfamily d, polypeptide 26 |
| Cyp4a14 | ■ | | ■ | cytochrome P450, family 4, subfamily a, polypeptide 14 |
| Cp | | ■ | ■ | ceruloplasmin |
| Slc7a12 | | ■ | ■ | solute carrier family 7 (cationic amino acid transporter, y+ system), member 12 |
| Kynu | | ■ | ■ | kynureninase |
| Slc6a6 | | ■ | ■ | solute carrier family 6 (neurotransmitter transporter, taurine), member 6 |
| Folh1 | | ■ | ■ | folate hydrolase 1 |

| DOWN REGULATED GENES | | | | |
|---|---|---|---|---|
| Gene | Female | HP | CR | Product name |
| Hsd17b11 | ■ | ■ | ■ | hydroxysteroid (17-beta) dehydrogenase 11 |
| Cd36 | ■ | ■ | ■ | CD36 molecule |
| Acsm3 | ■ | ■ | ■ | acyl-CoA synthetase medium-chain family member 3 |
| Agps | ■ | ■ | ■ | alkylglycerone phosphate synthase |
| Mpv17l | ■ | ■ | ■ | Mpv17 transgene, kidney disease mutant-like |
| Ldhd | ■ | ■ | ■ | lactate dehydrogenase D |
| Cyp7b1 | ■ | ■ | | cytochrome P450, family 7, subfamily b, polypeptide 1 |
| Myo5a | ■ | ■ | | myosin VA |
| Slc22a19 | ■ | ■ | | solute carrier family 22 (organic anion transporter), member 19 |
| Kif20b | ■ | ■ | | kinesin family member 20B |
| Slco1a1 | ■ | ■ | | solute carrier organic anion transporter family, member 1a1 |
| Pzp | ■ | ■ | | PZP, alpha-2-macroglobulin like |
| B4galt5 | ■ | ■ | | UDP-Gal:betaGlcNAc beta 1,4-galactosyltransferase, polypeptide 5 |
| Slc22a7 | ■ | ■ | | solute carrier family 22 (organic anion transporter), member 7 |
| Tc2n | ■ | ■ | | tandem C2 domains, nuclear |
| Slc6a15 | ■ | ■ | | solute carrier family 6 (neurotransmitter transporter), member 15 |
| Gclc | ■ | ■ | | glutamate-cysteine ligase, catalytic subunit |
| Hsd11b1 | ■ | ■ | | hydroxysteroid 11-beta dehydrogenase 1 |
| Lpl | ■ | | ■ | lipoprotein lipase |
| Kap | ■ | | ■ | kidney androgen regulated protein |
| Slc22a28 | ■ | | ■ | solute carrier family 22, member 28 |
| Slc17a3 | ■ | | ■ | solute carrier family 17 (sodium phosphate), member 3 |
| Pank1 | | ■ | ■ | pantothenate kinase 1 |
| Amacr | | ■ | ■ | alpha-methylacyl-CoA racemase |

**Supplementary Figure S21. Shared genes altered in Pax2 and Pax8 mutant S3 proximal tubule also differentially expressed in other mechanisms of IRI protection.** Genes included were differentially expressed in Mutant S3 compared with healthy S3 and at least 2 of 3 data sets: female sex[41] (female), hypoxic preconditioning[39] (HP), and caloric restriction[39] (CR) as indicated.

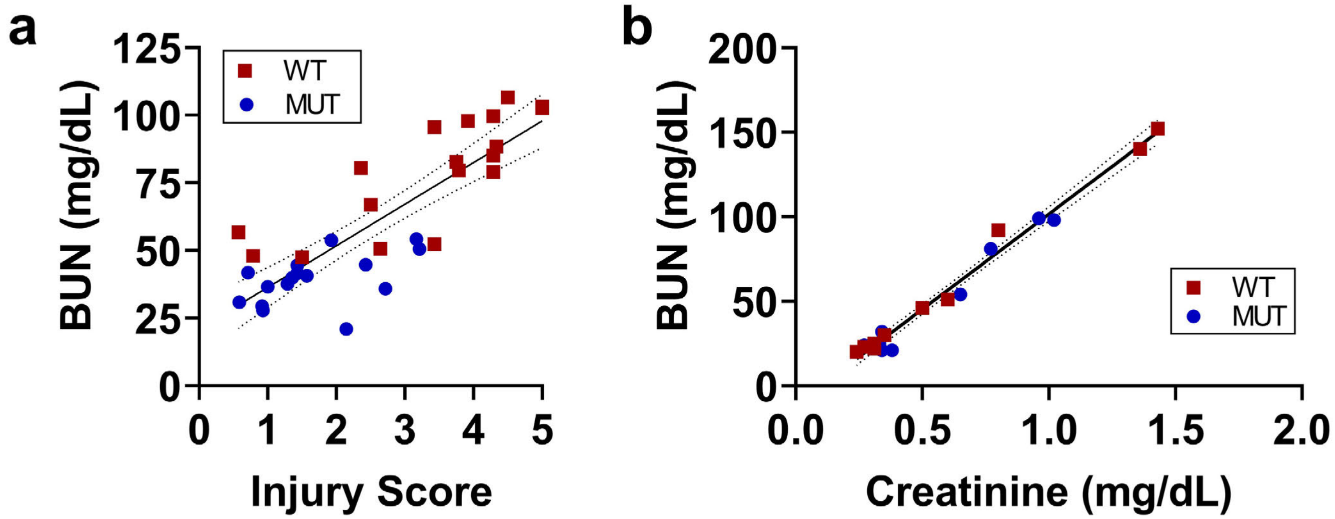


**Supplementary Figure S22. BUN correlates with histologic injury and serum creatinine.** (**a**) Terminal BUN measurements were highly correlated with histologic injury score in mutant and WT mice (Pearson r = 0.84, $P < 0.0001$). (**b**) BUN was tightly correlated with serum creatinine in terminal serum samples from injured and uninjured mutant and WT mice (Pearson r = 0.99, $P < 0.0001$).

**SUPPLEMENTARY METHODS:**

Animals. All studies involving mice were approved by the Institutional Animal Care & Use Committee (IACUC) at the University of Michigan and performed in accordance with the NIH Guide for the Care and Use of Laboratory Animals and ARRIVE guidelines. Phosphoenolpyruvate carboxykinase (PEPCK) Cre animals were obtained from Volker H. Haase[24]. Mice containing a GFP Cre reporter were purchased (Gt(Rosa)26Sor$^{tm4(ACTB\text{-}tdTomato,\text{-}EGFP)luo}$, Jackson Labs, Stock No. 007676). Mice with floxed Pax2 and Pax8 alleles were derived from the University of California, Davis Knockout Mouse Project repository (clone EPD0847_2_B03) and from Dr. M. D Felice[54], respectively, as used previously[21]. Strains used in these studies were derived from a common breeding pair of one PEPCK Cre$^{+/y}$, GFP reporter$^{+/+}$ (where + indicates the presence of the transgene allele) animal and one Pax2$^{fl/fl}$, Pax8$^{fl/fl}$ animal (where fl indicates the presence of a mutant allele with loxP sites flanking DNA binding exons in each gene). Offspring were bred to homozygosity to generate colonies used for subsequent studies. Except where noted, all animals were hemizygous for PEPCK Cre and homozygous for the GFP reporter. Only male animals were used to provide consistency with the prior IRI literature which largely uses male mice. Wild type (WT) strains contained Pax2$^{wt/wt}$ and Pax8$^{wt/wt}$ alleles and mutants (MUT) contained Pax2$^{fl/fl}$ and Pax8$^{fl/fl}$ alleles. Complete genetics for each strain is shown in Supplementary Figure S1. Animals with alternative genetics were used in some studies with Pax2$^{fl/fl}$ and Pax8$^{fl/fl}$ present in all animals where absence or presence of PEPCK Cre determined WT control and mutant strains, respectively (Supplementary Figure S8). PCR primers used for genotyping are provided in Supplementary Table S1.

We initially targeted 14-16 animals per group per timepoint for post-IRI analyses based on a meaningful effect size of 30%, standard deviation of 30% (typical for IRI[55]), $\alpha = 0.05$, and 80% power. The actual number of animals used for each experiment varied with breeding productivity at the time of surgery and revised estimates for effect size and variance. Animals were used between 11 and 13 weeks of age. Cage locations were randomly assigned in the housing facility. Animals were randomly ordered for surgery or euthanasia, but operators were aware of the set of animal genotypes included each day. Genotypes were unblinded for subsequent analyses after euthanasia.

Ischemia-reperfusion injury (IRI). Two surgical models were used. The first model resulted in severe unilateral IRI, modeling AKI-to-CKD transition. Age-matched 12-week-old male (range 11-13 weeks) mice were anesthetized with isoflurane. The retroperitoneum was accessed by a small incision on the left flank. The kidney was retracted gently, and fat was dissected from the pedicle. The renal artery and vein were clamped with a Schwartz clip (Roboz Surgical, RS-5459). After 30.0 min, the clamp was removed and reperfusion of the kidney was confirmed. The incisions were closed in layers. Body temperature was maintained at 38 ºC throughout the procedure using an infrared homeothermic control system (Kent Scientific, RightTemp Jr). In this model, the contralateral kidney was not manipulated. The second model resulted in moderate injury and included a pre-injury contralateral nephrectomy allowing serum-based measurements of kidney function. These surgeries were performed as above except that prior to clamping, the right kidney was exposed, ligated with non-resorbable suture, excised, and processed for additional analysis (see below). Clamp time was reduced to 25.0 min and homeothermic warming was set to 37 ºC. In both models, intraperitoneal (IP) warm sterile saline (20 µl/g) was provided after wound closure to compensate for fluid loss during surgery. Analgesia with carprofen (5 µg/g IP) was provided for 48 h post-operatively (exactly 2 doses, immediately and 24h after induction). Animals with surgical complications were excluded from analysis; this criterion was determined *a priori*. Animals were euthanized using carbon dioxide at times indicated.

For 6h and 2d (with contralateral nephrectomy) and 14d (without contralateral nephrectomy), there was no unexplained mortality owing to the short duration of follow up and preserved renal function provided by the intact contralateral kidney, respectively. For 28d (with contralateral nephrectomy), 5 of 20 (25%) and 2 of 18 (11%) animals died or were euthanized prematurely for WT and MUT animals respectively. These rates were not significantly different ($P = 0.41$, Fisher's exact test). Of the deaths, 2 of 5 and 2 of 2 were euthanized due to wound infections (WT and MUT respectively) with the balance found dead or moribund (then immediately euthanized) after the 7d perioperative monitoring period.

Sample preparation. Kidneys obtained from contralateral nephrectomy during IRI surgery were divided sagittally on ice. For one half, a small portion of one pole was removed to extract DNA and the remaining tissue was fixed in 4% formaldehyde. The remaining half kidney was quartered and frozen. For terminal sample collections, blood was obtained immediately after death followed by perfusion with ice-cold phosphate buffered saline (PBS) via the left ventricle. Kidney tissue was explanted and divided sagittally. Half was fixed overnight (12-18h) in 4% paraformaldehyde in PBS. Remaining tissue was quartered and flash frozen. Fixed tissue was dehydrated and embedded in paraffin. Multiple samples were embedded in each paraffin block. Sections (~5 µm thickness) from 1-4 blocks were mounted per slide. Each slide, therefore, contained samples from several animals. Slides were stored at 4ºC, desiccated, under nitrogen. Block location, orientation, and section shape were used to track and confirm the identity of each sample on the slide.

Immunofluorescence staining. Sections were deparaffinized and rehydrated using a graded series of xylene-ethanol-water solutions. Slides were treated with either citrate- or tris-based antigen unmasking solution (Vector Laboratories, depending on primary antibodies used) in a pressure cooker for 10 min. Samples were then permeabilized with 0.5% Triton-X 100 in PBS, blocked with 20 mg/ml bovine serum albumin, and incubated overnight with primary antibodies (Supplementary Table S2) in 0.1% Tween-20 in PBS (PBS-T). Sections were then extensively rinsed with PBS-T and incubated with species-specific fluorophore-conjugated secondary antibodies in PBS-T for 2 h at room temperature, rinsed with PBS-T, then PBS, then water, and quickly dried. Hydrophobic pen (which interferes with automated slide scanning) was removed by immersing the slides in xylene for 60 s. Xylene was then quickly evaporated under vacuum and the slides rehydrated in PBS. Samples were then treated with the TrueVIEW autofluorescence quenching kit (Vector labs). Samples were counterstained with 1 µg/ml DAPI. Samples were mounted with Prolong Diamond mounting medium (Invitrogen), allowed to cure overnight at room temperature protected from light, and stored at 4ºC protected from light until scanning. We utilized two methods for co-stains using same-species primary antibodies. In the first, the primary antibody was labeled with the FlexAble antibody labeling kit (Proteintech) and incubated with the sample for 2-4h following primary staining. For these samples, quenching and xylene steps were omitted due to incompatibility. In the second method, we repeated the indirect staining method after completing the primary stain but decreased incubation times to 2-3h. In this method, additional control sections were also included which omitted the primary antibody only from the second round of staining. These controls tested for secondary antibody cross reactivity with the primary stain, which was minimal. Both methods resulted in lower staining sensitivity compared with standard indirect immunofluorescence. Antibody suppliers and dilutions are listed in Supplementary Table S2.

Blood urea nitrogen (BUN) and creatinine measurement. BUN measurements were performed using the QuantiChrom Urea Assay Kit (BioAssay Systems). Creatinine was measured using a clinical bioanalyzer. Because BUN correlated with histologic injury score (see below) and creatinine across strains and degrees of injury (Supplementary Figure S22), we utilized BUN measurements as a surrogate for kidney function in most studies.

Image processing: Stained slides were scanned at a resolution of 0.5 µm/pixel and processed as described and extensively characterized previously[27] to generate segmentations of positive staining over the entire cortex and outer stripe of the outer medulla (OSOM) for each image. One sagittal section was analyzed per animal. Images with obvious artifacts (inconsistent or weak staining, tissue folds, etc) resulting in spurious segmentation results were excluded from analysis. U-Net[52] segmentation models were trained to identify whole cells, including the nucleus, not just strongly staining portions. For single stains, analysis of single channel segmentations was reported. However, to assess co-expression, tissue cytometry was performed. Each DAPI stained nucleus (identified by U-Net segmentation[27], typically 100-200,000 nuclei per section) was assigned to a positive or negative status based on its overlap with other target segmentations (for example, segmentations for Pax8 or GFP). Overlaps were considered positive if greater than 25% or 80% for nuclear and membrane-associated stains, respectively. Example tissue cytometry results are shown in Supplementary Figure S2, Supplementary Figure S5, Supplementary Figure S7, and Supplementary Figure S9. Data presented represents the average metric over the entire cortex + OSOM for each animal.

Injury scoring. Sagittal hematoxylin and eosin-stained sections were scanned at a resolution of 0.5 µm/pixel. Each section was then de-identified, randomized, and blurred followed by placement of 6-10, non-overlapping, 1 mm diameter circular ROIs around the cortex near the corticomedullary junction (as many as fit). Full resolution ROIs were extracted from the original image. Blinded ROIs were then randomly assigned to batches for scoring. Each batch contained 1 ROI from each sample. A duplicated but randomly rotated injured ROI was also included for each injured kidney to quantify scorer consistency. Image order in each batch was randomized. 6-8 batches were scored. The average of each sample's scored ROIs was reported. Acute injury (less than 4 day from IRI) was scored by calculating the percentage of tubules that display cell necrosis, loss of the brush border, cast formation, or tubular dilatation as described by Chen *et al*[56] using the cutoffs: 0: No injury, 1: 1-10%, 2: 11-25%, 3: 26-45%,4: 46-75%, 5: 76-100%. Chronic injury (greater than 4 day from IRI) was scored by calculating the percentage of tubules in the corticomedullary junction that display any matrix-rich expansion of the interstitium with infiltrating interstitial cells, chronic tubular injury (intratubular casts, atrophy of tubular cells with flattened/simplified tubular epithelial cells with thickened tubular basement membranes) as described by Borza, *et al*[55] using the cutoffs: 0: No injury, 1: 1-25%, 2: 26-50%, 3: 51-75%,4: 76-100%. Intra-scorer correlation was > 0.94.

Western blot. Frozen kidney samples were thawed on ice, minced with a razor blade, thoroughly mixed, and divided into equal portions for protein and RNA isolation. The portion of this tissue assigned to protein isolations was then homogenized in 15 µl/mg of radioimmunoprecipitation assay (RIPA) buffer with protease inhibitors (Santa Cruz Biotechnology) and 5 mM sodium fluoride with a handheld homogenizer on ice. The homogenate was then incubated on a rotor for 2 h at 4 ºC. Insoluble material was collected by centrifugation at 17,000×g for 10 min at 4 ºC. Protein concentration in the supernatant was determined using the bicinchoninic acid (BCA) assay (Thermo). Because more samples were available than can fit on a standard gel, we subsampled by matching injury severity score ranks (i.e. top 6-7 samples from each strain). Samples were then mixed with 2X sodium dodecyl sulfate (SDS) loading buffer and heated at 95ºC for 5 min prior to use. For each sample, 30 µg of protein was loaded in 4–15% SDS-polyacrylamide gel electrophoresis (PAGE) gels (Bio-Rad) followed by electrophoresis for 1 to 1.5 h. Protein was transferred to a polyvinylidene fluoride (PVDF) membrane (Millipore Sigma). Membranes were blocked with dehydrated milk and incubated with primary antibodies (Supplementary Table S2) overnight at 4°C. After rinsing, the membrane was incubated with an HRP-labeled secondary antibody. Signal was detected using enhanced chemiluminescence (ECL) reagent (Thermo) and autoradiography film.

RT-PCR. Minced kidney tissue (prepared above) was combined with 0.8 ml of ice cold TRIzol LS (Thermo) and homogenized on ice and RNA extracted per manufacturer protocol. RNA content was quantified using a Nanodrop spectrophotometer (Thermo). Reverse transcription was performed using the HighCapacity cDNA Reverse Transcription Kit (Thermo) per manufacturer instructions using random primers. qPCR was performed using a 7500 Real Time PCR System (Applied Biosystems, Thermo) with SYBR Green master mix according to manufacturer instructions with primers listed in Supplementary Table S3. For analysis of mutant and wild type Pax2 and Pax8 transcripts, PCR was performed in a standard thermocycler and visualized using agarose gel electrophoresis.

Single nucleus preparation and snRNA seq. Flash frozen kidney samples were stored at -80 °C prior to processing. For nucleus isolation, all reactions were carried out at 4°C. The specimens were thawed and minced thoroughly for one minute in NP40 lysis buffer (0.1% NP40, 10 mM Tris pH 7.5, 146 mM NaCl, 1 mM CaCl2, 21 mM MgCl2, and 40 U/mL RNAse inhibitor). Next, the suspension was further homogenized in 1 mL lysis buffer using a Dounce homogenizer (Kimble, cat:885300-0002) with 20 strokes each using pestle A and pestle B. Subsequently, 4 mL of wash buffer (10 mM Tris, 146 mM NaCl, 1 mM CaCl2, 21 mM MgCl2, 1% BSA [7.5% Fraction V], and 40 U/mL RNAse inhibitor) was added, filtered through a 30 µm strainer (Milteni), and centrifuged for 10 min at 200×g at 4 °C. The supernatant with debris was carefully removed using a glass Pasteur pipette and house vacuum, and the pellet was then resuspended in 5 mL wash buffer, filtered through a 5 µm strainer (Pluriselect), and centrifuged again. After removing the supernatant, the pelleted nuclei were resuspended in Parse nuclei buffer, counted, fixed, permeabilized, neutralized, and frozen at -80°C, following the Evercode fixation user manual V2.0.1 (Parse Biosciences) for nuclei. We applied the Split-seq snRNAseq method using two wells per sample (10,000 nuclei per well) of a 96-well "Evercode™ WT Mega v2" plate (capacity of ~1 million nuclei, Parse Biosciences) to generate single nuclei data for a total of eight samples. Barcoding (four rounds) was carried out according to the "Single Cell Whole Transcriptome Kit User Manual" (Parse Biosciences), followed by library generation and sequencing on an Illumina NovaSeq6000 platform (paired end) at a sequencing depth of 20K per nucleus, which was executed at the Advanced Genomics Core of the University of Michigan. The resulting FASTQ files were analyzed using the Parse Biosciences data analysis pipeline (v1.0.2p) which demultiplexes reads and assigns cells and samples. Reads were aligned to the mouse genome (GRCm38 mm10) with the addition of custom sequences for mT/mG transcripts (both tdTomato and EGFP) and Cre which included 3' UTRs.

The resulting data matrices were read into the Seurat package[26] (v4.3.0) in R. Outputs from the "Evercode™ WT Mega v2" plate were provided as 8 separate libraries (each with a mix of transcriptomes from all samples). Each library was imported using the “ReadParseBio” function and merged together to create a single Seurat object for the experiment. Quality control (QC) cut-offs of 200-5000 features per nucleus and less than 5% mitochondrial gene reads were used to filter raw sequencing data. This resulted in 62,237 total transcriptomes. The resulting raw data matrix was scaled, normalized and processed in a preliminary fashion to generate 28 preliminary clusters (using 30 principal components with resolution 0.3). These clusters were aligned to previous snRNA seq data from whole mouse kidney[6] in order to estimate the number of nuclei likely to be doublets based on dual alignment and divergent feature histograms. Based on these criteria we estimated 5567 likely doublets. We then processed this preliminary data with DoubletFinder (v2.0.3)[57] and proceeded with only the subset of 57,670 transcriptomes classified as likely singlets.

The top 2000 variable features in the resulting cleaned data set were used to generate cell clusters (using 30 principle components, resolution 0.3). Clusters were visualized using uniform manifold approximation and projection (UMAP). Differentially expressed genes in each cluster were identified using the “FindAllMarkers” function in the Seurat package. Cluster annotations were assigned by using the top 50 differentially upregulated genes for each cluster from Kirita et al[6], a similar single nucleus whole kidney dataset, as markers for specific kidney cell types. The lists of cluster markers in our data were ranked by average log2 fold change and evaluated by gene set enrichment analysis (GSEA) using the fgsea package in R[58] against these cell type marker sets from Kirita et al. Only upregulated genes were used for annotating clusters. The matrix of normalized enrichment scores (presented as a heatmap in Supplementary Figure S11) were used to annotate clusters based on maximal enrichment. A dot plot of marker genes used to validate clusters in Kirita et al was replicated for our data (Supplementary Figure S11). The number of cells in each cluster from each sample was also measured to ensure clusters were not biased by outlier samples (Supplementary Figure S12). Comparisons between mutant and WT cells within each cluster were performed with “FindMarkers” in Seurat.

Cells annotated as proximal tubule using this approach were extracted, reclustered, and new marker genes determined as above. Annotations were defined using GSEA as above but against the differentially upregulated marker genes only in the proximal tubules of healthy and post IRI animals from Kirita et al[6]. Comparisons between healthy S3 and mutant S3 proximal tubule (Figure 6D) were obtained using the “FindMarkers” function. In some cases, the activities of collections of genes were assessed by computing the average Z-score of the expression of each of the set elements, per cell, using the computeZScore function of the singleCellTK (v2.8.0, DOI: 10.18129/B9.bioc.singleCellTK ) package in R. Pathway activity was assessed using PROGENy[29-31] within the decoupleR (v2.4.0)[53] package. Transcription factor activity was assessed using signatures defined by DoRothEA (v1.10.0)[32], limiting analysis only to signatures with “A” level confidence.

GSEA for other comparisons was performed using fgsea as above with other gene sets (for example, caloric restriction) and expression lists ranked by log2 fold change (including both up and down regulated genes).

Pseudotime ordering was performed using Slingshot[35]. For clarity, the “Mixed” proximal tubule cluster was excluded from the analysis reported because it did not have clear segment-specific markers associated with it, suggesting it was artifactual in origin. The number of cells in this cluster was small and composed of both genotypes equally. Analysis with this cluster included did not reveal any trajectories linking the S3 with S3-Mut clusters.

Proximal tubule purifications. 12-week-old mice were euthanized and flushed with ice cold PBS followed by 30 mg/ml Collagenase, Type 4 (Worthington Biochemical). Kidneys were explanted, decapsulated and minced finely with a razor on ice. Minced kidney was digested at 37 ºC for 12 min at 200 RPM in pre-oxygenated DMEM/F-12 with 0.25 mg/ml Liberase TL (Roche) followed by trituration of partially digested tissue. The digestion was quenched with 10% heat inactivated FBS and cooled to 4 ºC. The resulting solution was passed twice through 40 µm cell strainers. Cells were then collected by centrifugation and a small portion was set aside (pre-purification). The remaining cells were incubated with biotinylated lotus tetragonolobus lectin (LTL, Vector Laboratories) for 10 min at room temperature in PBS. Cells were washed then incubated with anti-biotin magnetic microbeads (Miltenyi Biotec) for 15 min at 4 ºC. The labeled cells then were separated using magnetic activated cell sorting (MACS) on LS columns according to manufacturer protocol (Miltenyi Biotec). Positive and negative fractions were collected for downstream analysis.

RNA from the resulting purified proximal tubule fractions was isolated from tissue samples using RNeasy kits (Qiagen) including optional DNase treatment. RNA was subjected to RT-PCR for marker genes from proximal tubule, loop of Henle, and distal tubule to confirm proximal tubule enrichment.

Analysis of RNA seq from purified proximal tubules. RNA from purified proximal tubules was processed for routine poly-A library preparation and subjected to 150 bp paired-end sequencing according to the manufacturer's protocol (Illumina NovaSeq) targeting 40 million reads per sample. Reads were aligned to the mouse genome (GRCm38 mm10) with the addition of custom sequences for mT/mG transcripts and Cre as in single nucleus sequencing data. Differential expression was determined using DESeq2 (v1.38.2)[59]. GSEA was performed using fgsea as with single nucleus expression data.

Statistics. Data are presented as mean and error bars represent standard deviation unless otherwise specified. Each data point represents the indicated measurement from a single animal. Statistical analysis was performed using the test indicated in R (version 4.2.2, transcriptomic analyses) or GraphPad Prism (version 8.0.0, all other analyses). Parametric tests were used for continuous or near continuous normally distributed data. Non-parametric tests were used for other data types.

**Supplementary Table S1. Genotyping PCR Primers**

| Target | Forward Primer | Reverse Primer | Product Size (bp) |
|---|---|---|---|
| Pax2 flox | ACCATAGACATTAGAGGTGCAGAGGG | AAAGTGAGGGAAGCGTAGAGAAGCC | ~600 (floxed)<br>~500 (WT) |
| Pax8 flox | GAAAGTTCGAGGGAAGGGAGATC | CAGTTCTTTCAGTGGTCCCTCC | ~400 (floxed)<br>~300 (WT) |
| Cre transgene | CGAGTGATGAGGTTCGCAAG | TGAGTGAACGAACCTGGTCG | ~375 |
| No Cre[a] | GAGCCTCTTAGAGCCCAAATG | GCACATGACTGCAACACACA | ~300 |
| mT/mG reporter | CTCTGCTGCCTCCTGGCTTCT | TCAATGGGCGGGGGTCGTT | ~350 |
| No mT/mG[a] | CTCTGCTGCCTCCTGGCTTCT | CGAGGCGGATCACAAGCAATA | ~260 |

[a]insertion site without transgene

**Supplementary Table S2. Antibodies**

| Target | Supplier | Product Number | Dilution |
|---|---|---|---|
| Aldh1a1 | Proteintech | 60171-1-IG | 400 (IF) |
| Cdh6 | in house[a] | | 1600 (IF) |
| Col1a1 | Cell Signaling Technology | 72026 | 2500 (WB) |
| Cp | Proteintech | 66156-1-IG | 400 (IF) |
| Gc | Proteintech | 16922-1-AP | 400 (IF) |
| GFP | Invitrogen | A10262 | 400 (IF) |
| Ki67 | Biolegend | 652402 | 200 (IF) |
| Kim1 | Biotechne | AF1817 | 200 (IF)<br>2500 (WB) |
| OAT-1 | Proteintech | 26574-1-AP | 200 (IF) |
| Pax2 | in house[b] | | 3200 (IF) |
| Pax2 | Proteintech | 29307-1-AP | 100 (IF[c]) |
| Pax8 | Proteintech | 10336-1-AP | 100 (IF[c])<br>400 (IF) |
| pH3 (Ser10) | Millipore Sigma | 06-570 | 1600 (IF) |
| Sox9 | Biotechne | AF3075 | 400 (IF) |
| Vcam1 | Abcam | ab134047 | 400 (IF)<br>4000 (WB) |
| Villin | Santa Cruz | SC-7672 | 200 (IF) |
| β-actin | Cell Signaling Technology | 4970 | 10000 (WB) |

IF: immunofluorescence; WB: Western blot

[a]Mah, S. P., Saueressig, H., Goulding, M., Kintner, C. & Dressler, G. R. Kidney Development in Cadherin-6 Mutants: Delayed Mesenchyme-to-Epithelial Conversion and Loss of Nephrons. Developmental Biology 223, 38-53 (2000). https://doi.org:10.1006/dbio.2000.9738 PMID - 10864459

[b]Dressler, G. R. & Douglass, E. C. Pax-2 is a DNA-binding protein expressed in embryonic kidney and Wilms tumor. Proc Natl Acad Sci U S A 89, 1179-1183 (1992). https://doi.org:10.1073/pnas.89.4.1179

[c]dilution used when conjugated with FlexAble linker (Proteintech)

**Supplementary Table S3. Quantitative PCR Primers**

| Gene | Forward Primer | Reverse Primer |
|---|---|---|
| Havcr1 | GCAATGCCCATCTTCTGCTT | CCACTCGACAACAATACAGACC |
| Hnf4a | ACACGTCCCCATCTGAAG | CTTCCTTCTTCATGCCAG |
| Krt20 | TGACAATGCTAAGCTGGCTG | GCCAGGTCCTTGTTCAGTTC |
| Lrp2 | CCCATCGAGTACAGGTGTGA | CCCTGCAGTCAACTTTGTGG |
| Pax2[a] | TCCCGATGGATATGCACTGC | AGGATCCCACTGGGTCGTTA |
| Pax8[a] | CTCAGTCTGTGAGCGACTCC | ATGGGGAGGTTGAATGGCTG |
| Slc12a1 | CTGGCCATCTTCATCACCAC | TGAGAAGTCGTAGCCCAACC |
| Slc12a3 | ACTGGAGAGGCATTGATGGT | GCAAGGTAGGAGATGGTGGT |
| Vcam1 | AGTCTGTGGATGGCTCGTAC | TAGAGTGCAAGGAGTTCGGG |
| β-actin | TGCGTGACATCAAAGAGAAG | GATGCCACAGGATTCCATA |

[a]primers span DNA binding domain exons as shown in Supplementary Figure S1 yielding 2 products, one each for mutant and WT transcripts
